# Distinct Perception Mechanisms of BACH1 Quaternary Structure Degrons by Two F-box Proteins under Oxidative Stress

**DOI:** 10.1101/2024.06.03.594717

**Authors:** Shiyun Cao, Huigang Shi, Sheena Faye Garcia, Yuki Kito, Hui Shi, Hailey V. Goldberg, Jackeline Ponce, Beatrix Ueberheide, Luca Lignitto, Michele Pagano, Ning Zheng

**Affiliations:** Department of Pharmacology, Box 357280, University of Washington, Seattle, WA, USA; Howard Hughes Medical Institute, University of Washington, Seattle, WA, USA; Department of Biochemistry and Molecular Pharmacology; Laura and Isaac Perlmutter Cancer Center, New York University Grossman School of Medicine, New York, NY 10016, USA; Proteomics Laboratory, Division of Advanced Research Technologies, New York University Grossman School of Medicine, New York, NY 10016, USA; Cancer Research Center of Marseille (CRCM), CNRS, Aix Marseille Univ, INSERM, Institut Paoli-Calmettes, Marseille, France; Howard Hughes Medical Institute, New York University Grossman School of Medicine, New York, NY 10016, USA

## Abstract

The transcription factor BACH1 regulates heme homeostasis and oxidative stress responses and promotes cancer metastasis upon aberrant accumulation. Its stability is controlled by two F-box protein ubiquitin ligases, FBXO22 and FBXL17. Here we show that the homodimeric BTB domain of BACH1 functions as a previously undescribed quaternary structure degron, which is deciphered by the two F-box proteins via distinct mechanisms. After BACH1 is released from chromatin by heme, FBXO22 asymmetrically recognizes a cross-protomer interface of the intact BACH1 BTB dimer, which is otherwise masked by the co-repressor NCOR1. If the BACH1 BTB dimer escapes the surveillance by FBXO22 due to oxidative modifications, its quaternary structure integrity is probed by a pair of FBXL17, which simultaneously engage and remodel the two BTB protomers into E3-bound monomers for ubiquitination. By unveiling the multifaceted regulatory mechanisms of BACH1 stability, our studies highlight the abilities of ubiquitin ligases to decode high-order protein assemblies and reveal therapeutic opportunities to block cancer invasion via compound-induced BACH1 destabilization.

## INTRODUCTION

Oxidative stress is the imbalance between endogenous and exogenous pro-oxidant factors and the cellular antioxidant defense systems^1^. Acute and prolonged oxidative stress cause direct damages to cellular components and interfere with redox signaling, thereby contributing to many human pathologies such as cancer, diabetes, neurodegenerative disorders, and cardiovascular diseases^2,3^. To adapt to oxidative stress, cells have evolved various mechanisms to reduce the formation of reactive oxygen species (ROS), neutralize these toxic agents, and repair oxidative lesions. These antioxidant responses are temporally controlled by redox-sensitive transcription factors that modulate the expression of cytoprotective genes.

BACH1, also known as BTB (Broad-Complex, Tramtrack and Bric-à-brac) and CNC (cap ‘n’ collar) homology 1, is a master transcriptional repressor in heme homeostasis and antioxidant defense^4^. As a member of the basic leucine zipper (bZIP) protein family, BACH1 heterodimerizes with small musculoaponeurotic fibrosarcoma (sMAF) proteins and binds to the antioxidant response elements (AREs) of many antioxidant genes, suppressing their transcription under unstressed conditions^5^. One of the well-characterized BACH1 target genes is *HMOX1*, which encodes heme oxygenase 1 (HO-1), an enzyme with antioxidant activity owing to its ability to catabolize free heme as a cytotoxic pro-oxidant^6^. Other targets of BACH1 include cellular factors involved in iron homeostasis and redox regulation, such as ferritin heavy and light chains, NADP(H) quinone oxidoreductase 1, glutamate-cysteine ligase catalytic and modulator subunits, and thioredoxin reductase 1^7-9^. In response to heme overload or oxidative stress, the expression of these genes is induced by the transcriptional activator nuclear factor erythroid-derived 2-like 2 (NRF2), which competes with BACH1 for binding sMAF and accessing AREs^10-13^.

The antagonizing functions of BACH1 and NRF2 are dynamically balanced at their protein levels by redox signaling. A wealth of studies have revealed a critical role of the ubiquitin E3 ligase Kelch-like ECH-associated protein 1 (KEAP1) in regulating NRF2 activity in a redox-sensitive manner^14^. Under normal conditions, KEAP1 directly interacts with NRF2 and promotes its ubiquitination and degradation^15,16^. Under oxidative or electrophilic stress, covalent modification of highly reactive cysteine residues in KEAP1 impairs its E3 ligase activity, enabling newly synthesized NRF2 to rapidly accumulate and move to the nucleus to induce the expression of antioxidant genes^17,18^. Recent studies have identified two F-box proteins, FBXO22 and FBXL17, which mirror the function of KEAP1 to regulate the stability of BACH1 in response to elevated heme and ROS levels^19,20^. FBXO22 has been shown to ubiquitinate and promote heme-induced degradation of BACH1, which alleviates the repression of *HMOX1* expression. FBXL17, on the other hand, has been documented to contribute to BACH1 turnover when cells are challenged by hydrogen peroxide and hemin-induced oxidative stress.

FBXO22, FBXL17, and KEAP1 are substrate receptor subunits of the cullin-RING ubiquitin ligase (CRL) complexes, which are classified in five subfamilies: CRL1, CRL2, CRL3, CRL4A/B, and CRL5^21^. With more than 200 unique substrate receptors, the CRL E3 complexes regulate diverse cellular functions by ubiquitinating substrates with high specificity. As a substrate receptor of CRL3, KEAP1 features an N-terminal BTB domain that forms a stable homodimer, allowing the E3 ligase to simultaneously recognize two distinct degrons of NRF2 in the form of short linear motifs (SLiMs)^22^. How FBXO22 and FBXL17, two substrate receptors of CRL1, recognize BACH1 in response to oxidative stress, however, remains unknown. Importantly, the functional relationship between FBXO22 and FBXL17 in regulating BACH1 stability is poorly understood.

Beyond its role in the oxidative stress response pathway, BACH1 also targets genes involved in cell cycle control, apoptosis, and metabolism^7,23,24^. By positively regulating the expression of multiple pro-metastatic genes, BACH1 has been shown to promote cancer cell invasiveness^25-27^. In this study, we reveal the distinct and complementary mechanisms by which FBXO22 and FBXL17 perceive BACH1, whose degrons are encrypted in the quaternary structures of its dimeric BTB domain and functionalized by different forms of oxidative stress. By shedding light on the molecular basis of BACH1 regulation, our results not only reveal an unexpected substrate recognition mode of CRL E3s, but also inform potential therapeutic strategies targeting the transcription factor in human diseases.

## RESULTS

### Structure of the SCF^FBXO22-BACH1-BTB^ complex

BACH1 contains an N-terminal BTB domain, which mediates its homodimerization (Figure 1A). Previous studies have mapped the BACH1 BTB domain to be critical for FBXO22 interaction^20^. In a GST pull-down assay with purified recombinant proteins, we validated the direct binding between the BACH1 BTB domain and the full-length FBXO22 protein in complex with SKP1 (Figure 1B). We further confirmed the homodimeric nature of the BACH1-BTB domain by size exclusion chromatography coupled with multi-angle light scattering (SEC-MALS) analysis (Figure 1C). Despite its expected symmetric architecture, the BACH1 BTB dimer appeared to be associated with only a single copy of FBXO22-SKP1.

**Figure 1.**
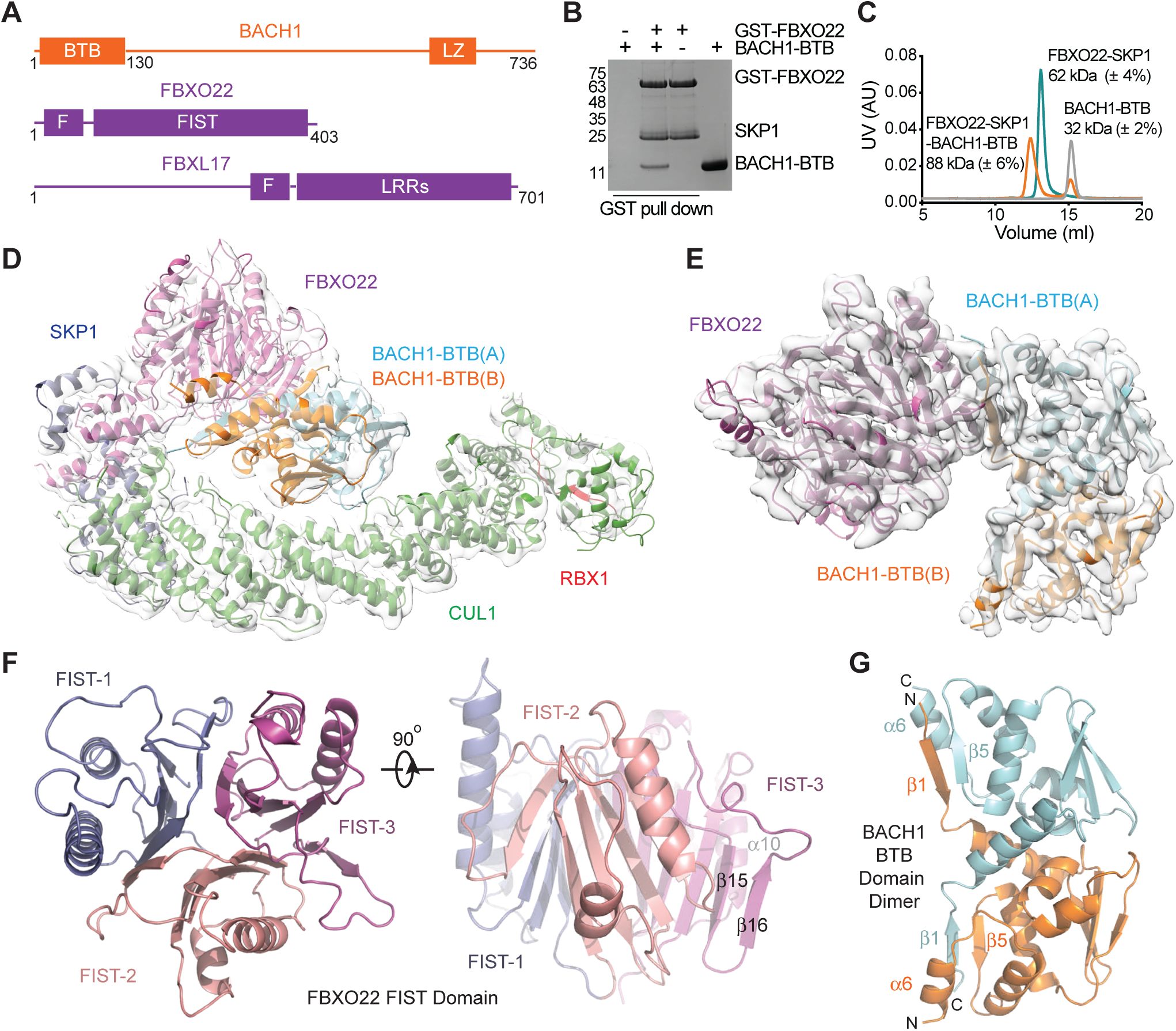
Overall structure of the FBXO22-BACH1-BTB complex. A. Domain composition of BACH1, FBXO22, and FBXL17. BTB: Broad-Complex, Tramtrack and Bric a brac; LZ: leucine zipper; F: F-box; FIST: F-box and intracellular signal transduction proteins; LRRs: leucine-rich repeats. **B.** BACH1-BTB pull down by GST-FBXO22. **C.** SEC-MALS analyses of BACH1-BTB, FBXO22-SKP1, and their complex with experimentally determined molecular weight shown next to each peak. Monomeric BACH1-BTB and FBXO22-SKP1 have a theoretical molecular weight of 17 kDa and 63 kDa, respectively. **D.** Structure model of SCF-FBXO22-BACH1-BTB fitted in the electron microscopy map. Subunits are labeled in different color. BACH1-BTB(A): light blue; BACH1-BTB(B): orange; FBXO22: purple; SKP1: dark blue; CUL1: green; RBX1: red. The RBX1 RING domain is not shown due to its flexibility. **E.** Electron microscopy map fit with the FBXO22-BACH1-BTB complex. **F.** Orthogonal views of the FBXO22 FIST domain. Three repeats (FIST-1, 2 and 3) are highlighted in different colors. **G.** Structure of the BACH1 BTB domain dimer with select secondary structure elements indicated.

To reveal the mechanism by which FBXO22 recognizes BACH1-BTB, we used cryo-electron microscopy (cryo-EM) to determine the structure of the SKP1-CUL1-RBX1-FBXO22 complex (also known as SCF^FBXO22^ or CRL1^FBXO22^) bound to the BACH1 BTB dimer at 3.9 Å resolution (Figure S1). The SCF^FBXO22^-BACH1 complex adopts a canonical SCF architecture^28^, in which the FBXO22-SKP1 substrate receptor module, together with the BACH1 substrate, is docked to the N-terminal half of the cullin scaffold (Figure 1D). Unique among F-box proteins, FBXO22 features a C-terminal FIST (F-box and intracellular signal transduction proteins) domain responsible for substrate recruitment (Figure 1A). Consistent with our SEC-MALS analysis, the F-box protein forms an asymmetric complex with the BACH1 BTB dimer at a 1:1 molar ratio (Figure 1E).

The FIST domain is found in eukaryotic F-box proteins and prokaryotic proteins involved in signal transduction^29^. It has been proposed to function as a sensory domain, possibly binding small molecular ligands, although its fold was previously uncharacterized. Our structure reveals that the FBXO22 FIST domain is constructed by three structural repeats, each consisting of a central four-stranded ꞵ-sheet sandwiched by two α-helices on one side and a ꞵ-hairpin loop on the other (Figure 1F, S2). The central ꞵ-sheet from each repeat packs against that of the other two repeats, giving rise to a compact globular fold with a pseudo three-fold symmetry. A Dali search identified two classes of unrelated metabolic enzymes, chorismatases and cyanuric acid hydrolases, with a similar overall fold, which could have evolved from monomers of the trimeric YjgF superfamily (Figure S2B)^30-32^. Different from the metabolic enzymes, the FIST domain of FBXO22 does not feature any obvious ligand-binding pocket. We name the three structural repeats of the FIST domain, FIST-1, FIST-2, and FIST-3. Noticeably, FIST-3 is distinguished from the other two repeats by having a longer ꞵ-hairpin loop, whose two ꞵ-strands, ꞵ15 and ꞵ16, perfectly align with the edge of the central ꞵ-sheet and extend it into a six-stranded sheet (Figure 1F, S2B). This FIST-3 ꞵ-hairpin, together with a nearby α-helix, α10, appear to play a critical role in recognizing the BACH1 BTB domain dimer.

### BACH1 quaternary structure degron

Similar to most BTB dimer structures, the BACH1 BTB domain dimer is characterized by a domain swapping architecture, in which the two protomers exchange their N-terminal ꞵ-strands^33,34^. In BACH1, this N-terminal ꞵ-strand (ꞵ1) of each protomer runs in antiparallel with the C-terminal α-helix (α6) of the opposite protomer, extending the dimer interface beyond the core BTB domain (Figure 1G). Remarkably, one of these cross-protomer ꞵ1-α6 pairs within the BACH1-BTB dimer makes up the entire interface with the F-box protein by clinging to the edge of FBXO22 FIST-3 (Figure 2A).

**Figure 2.**
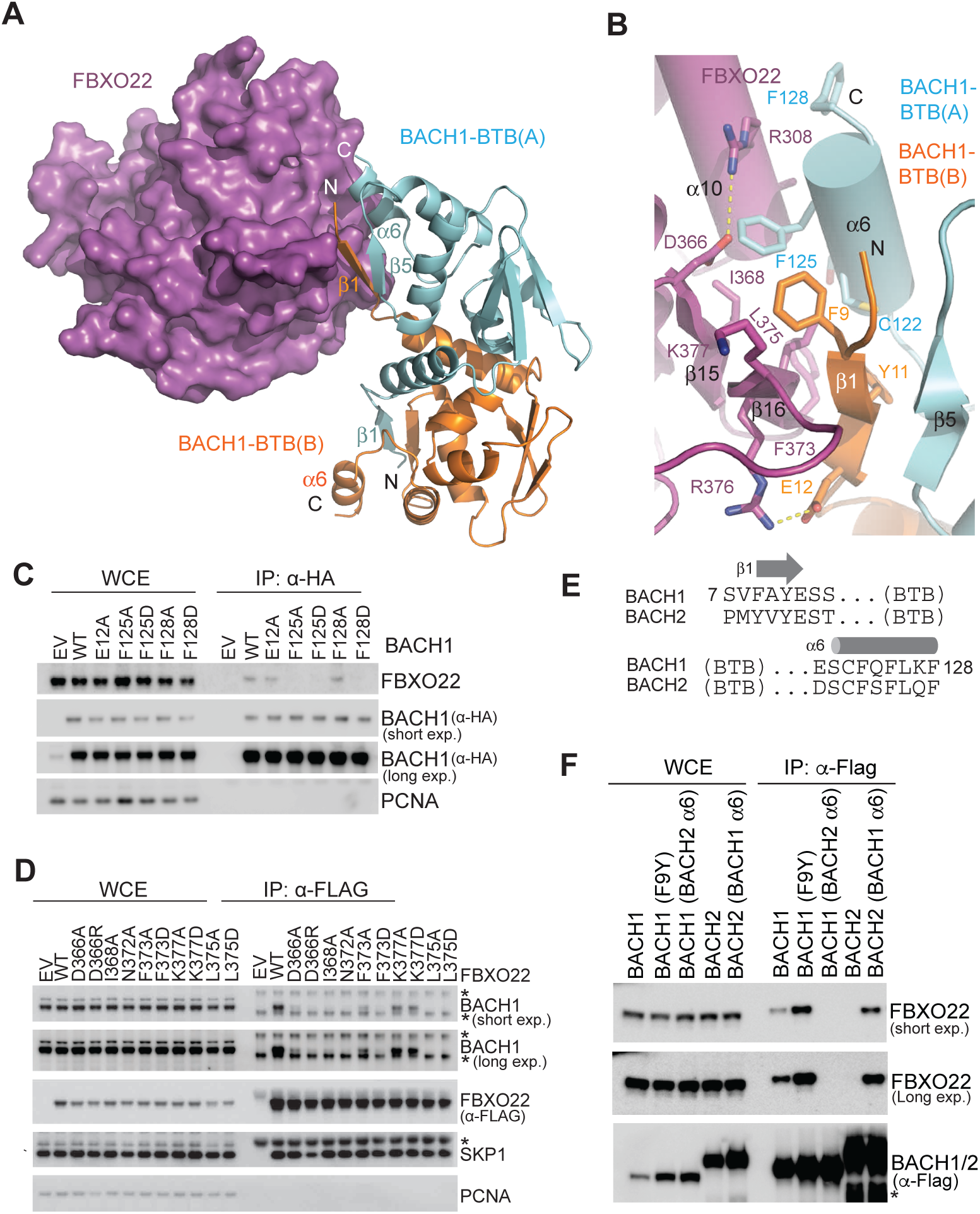
Recognition of the BACH1-BTB quaternary structure degron by FBXO22. A. Recognition of the BACH1 cross-protomer ꞵ1-α6 degron by FBXO22. BACH1-BTB(A) and BACH1-BTB(B) are shown is cartoons and colored in cyan and orange, respectively. FBXO22 is shown in purple surface representation. **B.** Close-up view of the FBXO22-BACH1-BTB interface with key residues shown in sticks. **C.** HEK293T cells were transfected with either empty vector (EV) or HA-tagged BACH1 wild type (WT) or indicated mutants for 48h. Cells were then treated with 2 μM MLN-4924 for 3h prior to HA-immunoprecipitation. Proteins were immunoblotted as indicated. **D.** HEK293T cells were transfected with either empty vector (EV) or 2xFLAG-tagged FBXO22 wild type (WT) or indicated mutants for 48h. Cells were then treated with 2 μM MLN-4924 for 3h prior to FLAG-immunoprecipitation. Proteins were immunoblotted as indicated. Non-specific protein bands are labeled with asterisks. **E.** Sequence alignment of the N- and C-terminal regions of human BACH1 and BACH2 BTB domains. **F.** HEK293T cells were transfected with either empty vector (EV) or 2xFLAG-tagged BACH1/BACH2 chimeras as indicated for 48h. Cells were then treated with 10 μM MG-132 for 3h prior to FLAG-immunoprecipitation. Proteins were immunoblotted as indicated. Non-specific protein bands are labeled with asterisks.

The tri-molecular interface between FBXO22 and the BACH1-BTB dimer is stabilized by a hybrid of polar and hydrophobic interactions. Specifically, the ꞵ1 strand of one BACH1-BTB chain (BTB-B), together with the ꞵ5 strand of the second BACH1 BTB chain, is juxtaposed with the ꞵ15-ꞵ16 hairpin of FBXO22 FIST-3, creating an intermolecular anti-parallel ꞵ-sheet supported by a network of backbone hydrogen bonds (Figure 2B). These secondary structure interactions are cemented by an adjacent hydrophobic core, which is nucleated by four amino acids from the BACH1-BTB ꞵ1-α6 pair (Phe9, Tyr11, Cys122, and Phe125) and three residues in the FBXO22 ꞵ15-ꞵ16 hairpin (Ile368, Phe373, and Leu375). Peripheral to the hydrophobic core, the FBXO22-BACH1-BTB dimer interface is further strengthened by two salt bridges and intermolecular van der Waal packing made by additional residues from the F-box protein (Asp366, Arg308, Lys377, and Arg376) and the two protomers of the substrate (Glu12 and Phe128).

In support of a critical role played by the N-terminal ꞵ1 strand of BACH1-BTB at the interface, previous studies have shown that individual alanine mutations of its two aromatic residues, Phe9 and Tyr11, are sufficient to abolish FBXO22-BACH1 interactions^20^. Similarly, replacing Phe125 in the BACH1 α6 helix with alanine or aspartate effectively abrogates the E3-substrate complex formation (Figure 2C). We conclude that both secondary structure elements of the BACH1-BTB ꞵ1-α6 pair, which is shaped by the quaternary structure of the BTB dimer, are necessary to define the functional degron. In a second set of mutational analyses, we validated that the majority of the FBXO22 residues at the tri-molecular interface are indispensable for binding BACH1, suggesting that the edge of the third FBXO22 FIST repeat represents the primary docking site for BACH1 (Figure 2D). Noticeably, although the symmetric BACH1 BTB domain dimer contains two cross-protomer ꞵ1-α6 pairs, two FBXO22 molecules cannot simultaneously recognize the two identical BACH1 degrons without a severe steric collision (Figure S3A). A single SCF^FBXO22^ E3 complex, therefore, appears responsible for ubiquitinating both protomers of a BACH1 dimer (Figure S3B).

Mammalian BACH1 has a close paralog, BACH2, which is primarily expressed in the brain and spleen with well-documented roles in regulating immune cell differentiation^35-37^. The BTB domains of the two BACH proteins, including their terminal regions, are overall 62% identical in sequence (Figure 2E). Their homodimer structures can be superimposed with a root mean square deviation (R.M.S.D.) of 0.6 Å (Figure S3C)^33,38^. Despite their sequence and structural similarity, only BACH1, but not BACH2, can be recognized by FBXO22 (Figure 2F). To identify key elements in BACH1 conferring this specificity, we made chimeric BACH proteins and tested their interactions with the F-box protein.

Interestingly, replacing Phe9 in the N-terminal ꞵ1 strand of BACH1 with its counterpart in BACH2 (Tyr12) had little effect on FBXO22 binding. By contrast, swapping the C-terminal α6 helices of the two BACH proteins effectively switched their ability to engage the E3 (Figure 2F). A close examination of the BACH2 BTB domain structure reveals a non-helical nature of its C-terminal region (Figure S3C) distinct from the α6 helix of BACH1. This structural difference provides a possible explanation for BACH2’s inability to bind FBXO22.

Together, these results reinforce the importance of the structural integrity of the ꞵ1-α6 degron of the BACH1 BTB dimer for FBXO22 recognition and underscore the capacity of the quaternary structure degron in conferring high E3-substrate selectivity, a previously undescribed characteristic of CRL substrates.

### Regulation of FBXO22-BACH1 interaction

Using BioLayer Interferometry (BLI), we determined the affinity of FBXO22 toward BACH1-BTB, which is ∼90 nM and on par with other CRL-substrate interactions (Figure 3A)^39^. Consistent with the lack of any obvious ligand-binding pocket in FBXO22 or BACH1-BTB, the binding between the two proteins is insensitive to heme (Figure 3B). Heme, therefore, does not appear to be physically required for the productive interaction between FBXO22 and BACH1 (Figure S3B) and most likely promotes BACH1 recognition by FBXO22 through an indirect mechanism. In previous studies, heme has been reported to block the DNA binding activity of BACH1-sMAF heterodimers independent of the BACH1 BTB domain^10^. This raises the possibility that the quaternary structure degron of BACH1 might not be accessible to FBXO22 until the transcription factor is released from the DNA.

**Figure 3.**
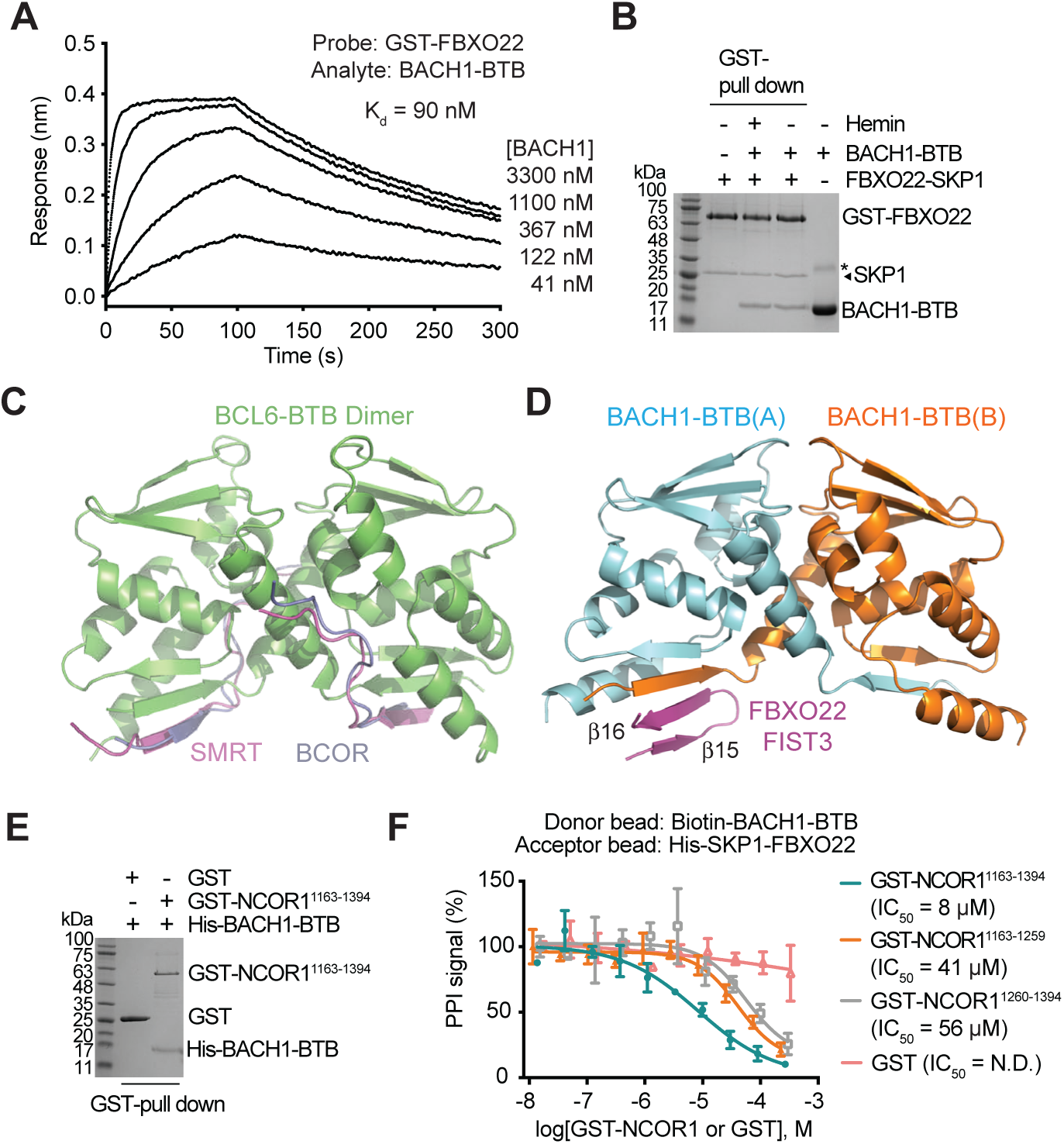
Regulation of FBXO22-BACH1 interaction. A. BioLayer Interferometry (BLI) measurements of FBXO22 and BACH1-BTB interaction. K_d-equilibrium_, dissociation constant. **B.** BACH1-BTB pull down by GST-FBXO22 in the presence and absence of 10 µM hemin. Non-specific protein bands are labeled with asterisks. **C.** Structure of BCL6-BTB dimer (green) in complex with the co-repressor SMRT (purple, PDB:1R2B) or BCOR (slate, PDB:3BIM). **D.** Structure of BACH1-BTB dimer (protomer A: light blue, protomer B: orange) in complex with FBXO22 ꞵ15-ꞵ16 hairpin (purple). **E.** BACH1-BTB pull down by GST-NCOR1. **F.** Inhibition of the FBXO22-BACH1-BTB interaction by NCOR1 fragments measured in AlphaScreen competition assays. IC50, half-maximum inhibitory concentration.

The BTB domain is widely found in CRL3 substrate receptors and transcription factors^34^. Besides mediating protein oligomerization, the BTB domain of select transcription repressors, such as PLZF and BCL6, has been reported to interact with co-repressors^40,41^. In the case of BCL6, distinct SLiMs of SMRT/NCOR2 and BCOR co-repressors have been shown to form the same interface with the BCL6 BTB domain dimer, which is highlighted by an inter-molecular ꞵ-sheet involving the N-terminal ꞵ1-strand of BCL6-BTB (Figure 3C)^42,43^. Such a binding mode is remarkably analogous to the FBXO22-BACH1-BTB interface (Figure 3D). Similar to PLZF and BCL6, BACH1 has been previously shown to be functionally associated with nuclear co-repressor NCOR1, a close paralog of SMRT/NCOR2^23^. This prompted us to test whether NCOR1 can compete with FBXO22 for binding the BTB domain of BACH1. A central region of NCOR1 has been previously identified to interact with the BTB domain of multiple functionally unrelated transcriptional co-repressors^40^. Sequence alignment and structure prediction analyses indicate that this region contains several potential SLiMs conserved among NCOR1 orthologs, including the one used by NCOR2 for binding BCL6-BTB (Figure S3D). When co-expressed and co-purified from *E. coli*, this ∼230 amino acid NCOR1 fragment was indeed sufficient to form a complex with the BACH1 BTB domain (Figure 3E). Importantly, the NCOR1 fragment was able to effectively block FBXO22-BACH1 binding in an AlphaScreen-based competition assay (Figure 3F), suggesting that the transcriptional co-repressor can occupy the quaternary structure degron of BACH1. Interestingly, when the N-terminal and C-terminal halves of the NCOR1 central fragment were individually tested, each sub-fragment was able to inhibit the interaction between FBXO22 and the BACH1 BTB dimer, albeit at a much lower potency (Figure 3F). Therefore, NCOR1 might employ more than one SLiM to simultaneously engage the two identical FBXO22-binding sites on BACH1-BTB to preclude the F-box protein. Together, these results imply that the NCOR1 co-repressor can shield the quaternary structure degron of chromatin-bound BACH1 from FBXO22 until the BACH1 is released from DNA by heme.

### Degradation of S-nitrosylated BACH1 by FBXO22 and FBXL17

FBXL17 is a leucine-rich repeats (LRRs)-containing F-box protein (Figure 1A) best known for its function in protein dimerization quality control^44^. Previous studies proposed that FBXL17 can recognize mismatched heterodimers of BTB proteins and sequester their monomeric BTB domains to ubiquitinate and degrade the non-functional ill-fated substrate proteins^45,46^. Under oxidative stress, how FBXL17 captures BACH1, presumably by recognizing its homodimeric BTB domain, remains unclear. We first investigated the functional relationship between FBXL17 and FBXO22 in regulating BACH1 stability in the context of BACH1 S-nitrosylation. Nitric oxide (NO) is a free radical in the cell that is either produced endogenously or exogenously^47^. By reacting with cysteine residues of selective targets, NO induces protein S-nitrosylation and further disulfide formation, which contribute to redox signaling that protects the cell against oxidative stress^48,49^. NOR3, an NO donor, has recently been shown to induce BACH1 degradation, an effect that can be augmented by a garlic-derived sulfur amino acid, S-1-propenylcysteine (S1PC)^50^. We first confirmed by biotin switch assay that BACH1 can be modified by two additional NO donors, S-nitrosoglutathione (GSNO) and spermine NONOate (sNONO) (Figure 4A). Using parental and FBXO22 knock-out HCT-116 cells, we verified that FBXO22 is responsible for destabilizing the majority of BACH1 at steady state and upon NO donor treatment (Figure 4B). By depleting FBXL17 in these cells, we further validated the role of FBXL17 in promoting NO donor-induced BACH1 degradation. The two F-box proteins, therefore, appear to play nonredundant roles in regulating BACH1 stability in stress responses.

**Figure 4.**
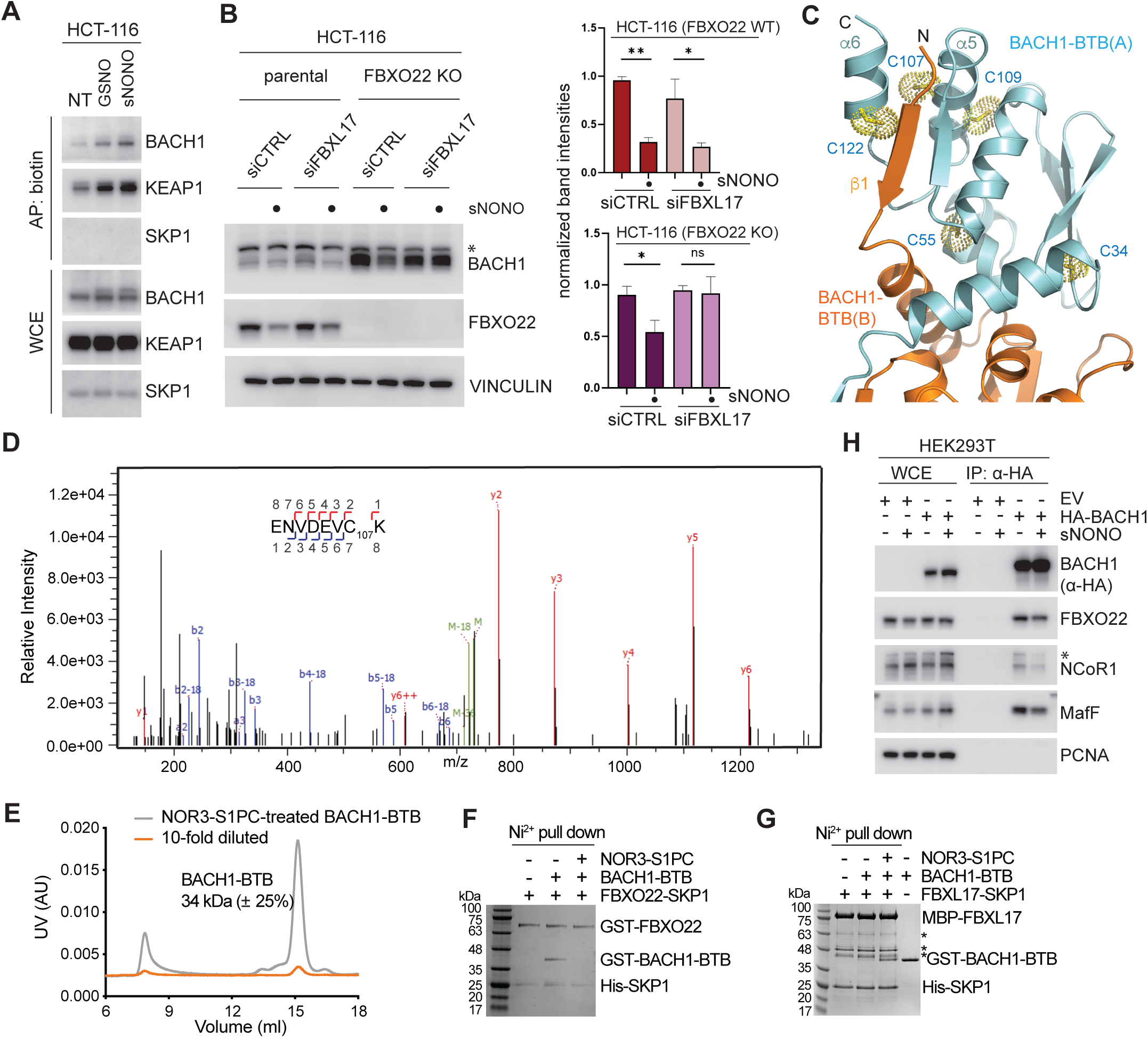
Degradation of S-nitrosylated BACH1 by FBXO22 and FBXL17. A. HCT-116 cells were treated with either DMSO (NT) or NO donors (GSNO and sNONO) for 6h before lysates were subjected to S-nitrosylation biotin switch. KEAP1 and SKP1 were included as positive and negative control, respectively^59,60^. Biotinylated proteins were affinity pulled down with streptavidin beads. Proteins were immunoblotted as indicated. **B.** *FBXO22* +/+ (parental) or *FBXO22* -/-(KO) HCT-116 cells were transfected with either non-targeting siRNA (siCTRL) or siRNA targeting FBXL17. Cells were treated with either DMSO (NT) or 100 μM sNONO for 6h as indicated. Proteins were immunoblotted as indicated. The graphs show the quantification of BACH1 protein levels. Values are presented as means ± SEM based on triplicates. **C.** Five cysteine residues in the BACH1-BTB domain with their side chains shown in dot spheres. **D.** An MS/MS fragmentation spectrum assigned to the S-nitrosylated peptide from BACH1-BTB containing Cys107 together with the assignments of the fragmentation series to the sequence of the modified peptide. **E.** SEC-MALS analyses of BACH1-BTB treated with NOR3-S1PC with experimentally determined molecular weight. The theoretical molecular weight of BACH1-BTB is 17 kDa for monomer or 34 kDa for dimer. **F.** Effects of BACH1-BTB S-nitrosylation on FBXO22 binding investigated by Ni^2+^ pull-down assay. **G.** Effects of BACH1-BTB S-nitrosylation on FBXL17 binding investigated by Ni^2+^ pull-down assay. Non-specific protein bands are labeled with asterisks. **H.** HEK293T cells were transfected with either empty vector (EV) or HA-tagged BACH1. Cells were treated with either DMSO (NT) or 100 μM sNONO prior to lysis. Lysates were subjected to immunoprecipitation with anti-HA beads. Proteins were immunoblotted as indicated.

The BTB domain of BACH1 contains five cysteine residues, two of which, Cys107 and Cys122, are located at the ꞵ1-α6 degron recognized by FBXO22 (Figure 4C). Consistent with a recent proteome-wide S-nitrosylation site mapping study^51^, mass spectrometry analysis of the recombinant BACH1 BTB domain treated with the NO donor NOR3-S1PC identified Cys107, but not Cys55, as an S-nitrosylation site (Figure 4D). Due to incomplete sequence coverage, whether Cys34, Cys109, and Cys122 could be modified by NO remained unclear. As a residue on the α5 helix of BACH1-BTB making direct contact with the α6 helix (Figure 4C), covalent modification of Cys107 is expected to perturb the local structure of the ꞵ1-α6 degron, thereby, impairing FBXO22 binding and possibly compromising the BACH1 BTB dimer interface. SEC-MALS analysis indicated that the NOR3-S1PC-treated BACH1 BTB domain still retains its homodimeric form in solution (Figure 4E). However, the NO donor-treated BACH1-BTB dimer can no longer interact with FBXO22 in a Ni^2+^ NTA pull-down assay (Figure 4F). By contrast, FBXL17 showed robust activity in binding the NOR3-S1PC-treated BACH1 BTB domain (Figure 4G). Interestingly, treatment with an NO donor appeared to reduce the binding of BACH1 to MafF and NCOR1 in the cell (Figure 4H), suggesting that a fraction of BACH1 was released from chromatin. This effect was accompanied with a decrease in BACH1 binding to FBXO22. These phenomena support the notion that S-nitrosylation impairs the ꞵ1-α6 degron of a sub-population of BACH1, thereby affecting its interaction with both FBXO22 and NCOR1. Taken together, these data strongly suggest that FBXL17 has been evolved to recognize the homodimeric BACH1 BTB domain when it is in a form incompatible with FBXO22 binding.

### FBXL17 preferentially remodels NO-modified BACH1-BTB dimer

From the single time point (30 mins) affinity pull-down assay, we noticed that FBXL17 preferentially interacts with the NO-modified BACH1 BTB domain over the native polypeptide (Figure 4G). To further dissect the activity of FBXL17 in BACH1-BTB recognition, we used BLI to monitor the kinetics of their interaction with or without treating BACH1-BTB with NOR3 and S1PC. In agreement with the affinity pull-down results, compound-treated BACH1-BTB interacted strongly with FBXL17 with an “apparent K_d_” of ∼110 nM, whereas the native BACH1-BTB protein only showed a detectable affinity towards the F-box protein with an “apparent K_d_” greater than 2.4 μM (Figure 5A, B). Noticeably, with compound-treated BACH1-BTB immobilized on the probe, the maximum binding signal appeared far from plateauing within the time window of the measurement, even though the amplitude of the association curves started to show saturation when the FBXL17 concentration reached the μM range (Figure 5A). The same phenomenon held true when we lengthened the time window to 10 minutes (Figure S4A). This feature hinted at a minimally biphasic nature of the interaction, which might involve complex rearrangement in addition to simple binding equilibrium.

**Figure 5.**
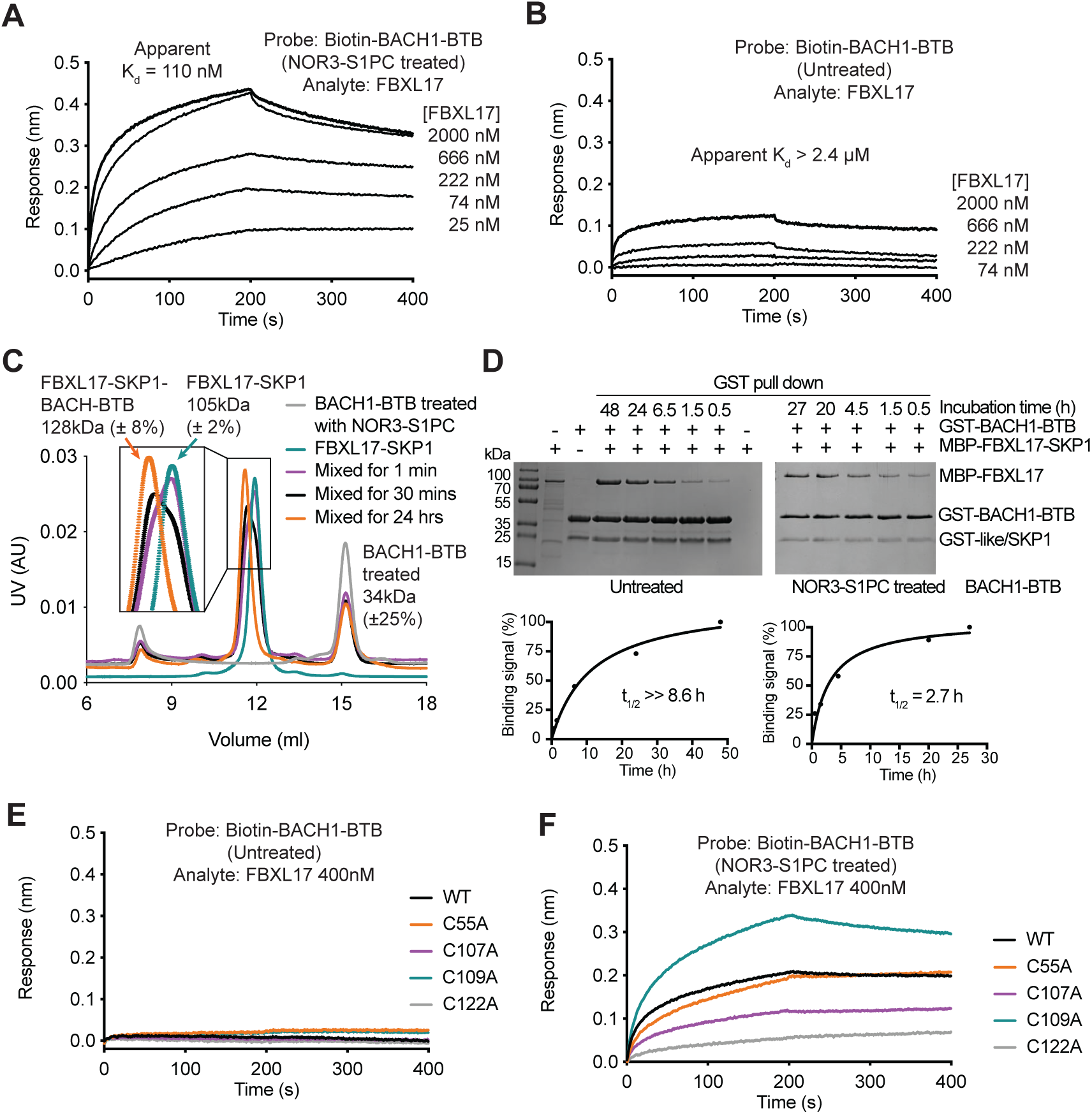
FBXL17 preferentially remodels S-nitrosylated BACH1-BTB. A. & B. BLI measurements of FBXL17 and BACH1-BTB interaction with and without NOR3-S1PC treatment. The apparent K_d-equilibrium_ values are indicated. **C.** SEC-MALS analyses of BACH1-BTB, FBXL17-SKP1, and their mixture at various incubation time with the experimentally determined molecular weights. The theoretical molecular weights of MBP-FBXL17-SKP1 and monomeric BACH1-BTB are 108 kDa and 17 kDa, respectively. **D.** The time-courses of the interactions between FBXL17 and BACH1-BTB with or without NOR3-S1PC treatment monitored by GST pull down. t1/2: time needed for 50% complex formation. **E. & F.** BLI measurements of the interaction between FBXL17 and BACH1-BTB (wild type and cysteine mutants) with or without NOR3-S1PC treatment. The effect of compound treatment was diminished by C107A and C122A but was enhanced by C109A.

To further characterize the interaction between FBXL17 and compound-treated BACH1-BTB, we used SEC-MALS to analyze samples of the two proteins mixed for different periods of time. In congruence with the reported activity of FBXL17 in forming stable complexes with monomeric BTB domains^45^, the F-box protein was found to bind a single copy of BACH1-BTB after the two proteins were mixed for 24 hours (Figure 5C).

Interestingly, it took a relatively long incubation time for such a stable 1:1 complex to form, even though BLI could instantaneously detect the interaction between FBXL17 and BACH1-BTB. Given that compound-treated BACH1-BTB remains a stable dimer even after being diluted to ∼3 μM (Figure 4E), we speculate that the biphasic binding curve detected by BLI reflects both the initial contact between FBXL17 and dimeric BACH1-BTB and the final complex formation between FBXL17 and monomeric BACH1-BTB after the BACH-BTB dimer is remodeled by the F-box protein.

To fully assess the binding kinetics of FBXL17 and BACH1-BTB, we returned to the affinity pull-down assay and monitored the interaction with a time window up to 1-2 days. As expected from our BLI and SEC-MALS results, it took about 2-3 hours for half of the maximal amount of FBXL17 to become stably associated with compound-treated BACH1-BTB under the tested experimental condition (Figure 5D). By contrast, the amount of FBXL17 captured by untreated BACH1-BTB did not saturate even after 48 hours. Such a strong preference of FBXL17 to capture BACH1-BTB monomers from the compound-treated dimer over the intact non-treated dimer supports the notion that NO treatment destabilizes BACH1 by promoting FBXL17 binding and subsequent ubiquitination.

Importantly, this effect of NO is attributable to the modification of two specific cysteine residues on the BACH1 BTB domain. Without impacting the resistance of native BACH1-BTB dimer to FBXL17 binding (Figure 5E), individual alanine mutations of Cys107 and Cys122, but not the other three cysteine residues, significantly reduced the ability of FBXL17 to release monomeric BACH-BTB from its NO donor-treated dimeric form (Figures 5F, S4B, S4C). This result allows us to map Cys122, an amino acid also residing within the ꞵ1-α6 degron, as another key residue sensitive to S-nitrosylation.

### Transient remodeling of BACH1-BTB dimer by a pair of CRL1^FBXL17^

To reveal the mechanism by which FBXL17 recognizes and remodels NO-modified homodimeric BACH1-BTB, we mixed NO donor-treated BACH1-BTB with the SCF^FBXL17^ complex for different periods of time and subjected the samples to cryo-EM analysis. 2D classification of the particles imaged at three incubation times (30 seconds, 10 minutes, and 24 hours) indicated that the predominant observable species were either in the form of free SCF^FBXL17^ or the E3 complex bound to the monomeric BACH1 BTB domain (hereafter referred to as mSCF^BTB^) (Figure S4D-F). Strikingly, a small fraction of the particles appeared to contain a pair of parallel SCF^FBXL17^ complexes with the FBXL17 LRR domains facing each other, likely sandwiching a BACH1-BTB dimer in the middle (Figure 6A). However, these minor classes, which possibly represent a transient intermediate species of the BACH1-BTB dimer remodeled by the E3 complex, had too few particles for high resolution 3D reconstruction.

**Figure 6.**
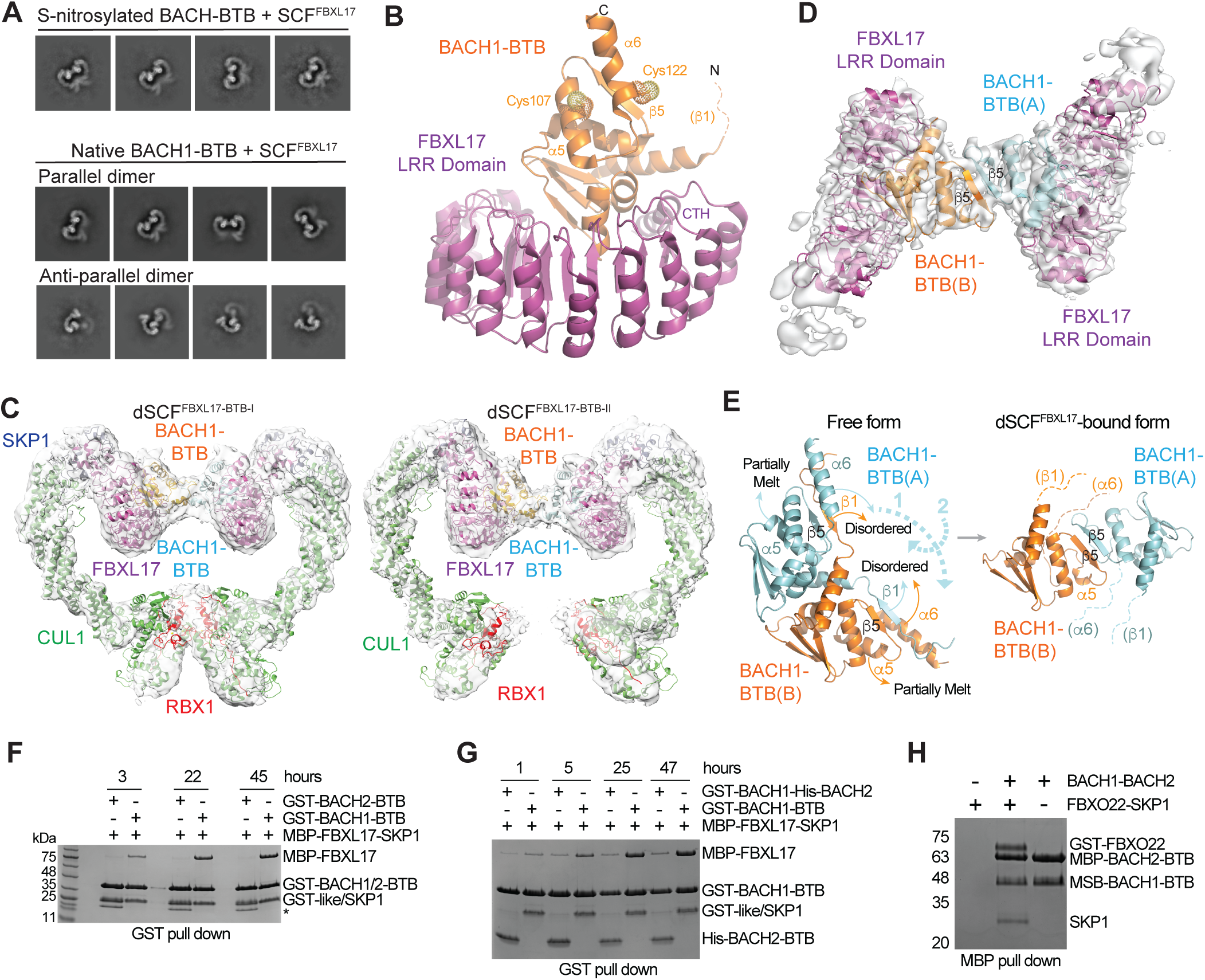
Structural analysis of BACH1-BTB remodeling by SCF^FBXL17^. A. Representative 2D classes of dimeric SCF-FBXL17-BACH1-BTB complexes found in the native and NOR3-S1PC-treated samples. **B.** Structure of FBXL17-LRR in complex with monomeric BACH1-BTB. Structurally disordered ꞵ1 strand of BACH1-BTB is shown in a dashed line. **C.** Two dimeric SCF-FBXL17-BACH1-BTB complexes fitted in their electron microscopy maps. Subunits are labeled in different color. BACH1-BTB (protomer A): light blue; BACH1-BTB (protomer B): orange; FBXL17: purple; SKP1: dark blue; CUL1: green; RBX1: red. **D.** Close-up views of BACH1-BTB dimer interface in the dimeric FBXL17-BACH1 complex. Local refinement-improved electron microscopy map is fitted with a BACH1-BTB dimer and two copies of FBXL17-LRRs. **E.** Conformational changes of the BACH1-BTB dimer during the transition from the free to the dimeric SCF-FBXL17-bound form. Two arrowed dashed lines indicate the two-step rotations of the protomer A (cyan) of BACH1-BTB. **F.** FBXL17 pull-down by GST-tagged BACH1-BTB but not BACH2-BTB. Non-specific protein band is labeled with an asterisk. **G.** A comparison of the efficiency of BACH1-BTB homodimer and BACH1-BTB-BACH2-BTB heterodimer remodeling by FBXL17 as monitored by the time course of GST-BACH1-BTB-MBP-FBXL17 association. **H.** Binding of the BACH1-BTB-BACH2-BTB heterodimer to FBXO22 by MBP pull-down assay.

To overcome the challenge of resolving these presumably unstable intermediates, we turned to the native BACH1-BTB sample, which is still susceptible to reconfiguration by FBXL17, except at a much slower rate. We reasoned that the higher resistance of the native BACH1-BTB sample to FBXL17-mediated monomerization might offer an experimental window to capture the intermediate steps of the reaction. Remarkably, cryo-EM analysis of a sample of SCF^FBXL17^ incubated with native BACH1-BTB unveiled four distinct species, free SCF^FBXL17^, mSCF^BTB^, and two dimeric SCF^FBXL17-BACH1-BTB^ super-assemblies (hereafter referred to as dSCF^BTB-I^ and dSCF^BTB-II^) (Figure S5). The structure of the mSCF^BTB^ complex resolved at 3.2 Å resolution is highly analogous to the previously reported crystal structure of FBXL17 in complex with the monomeric KEAP1 BTB domain^45^, except that no clear density is present for the ꞵ1 strand of the BACH1 BTB domain (Figure 6B). In both structures, the canonical dimer interface of the BTB domains is blocked by the FBXL17 C-terminal α-helix (CTH). Importantly, both Cys107 and Cys122 of BAHC1-BTB are solvent-exposed and far away from the BACH1-FBXL17 interface, indicating that they are not directly recognized by the F-box protein. Modification of these two residues in the NO donor-treated BACH1 sample, therefore, likely accelerated FBXL17 binding by destabilizing the BACH1-BTB dimer.

Although the 3D reconstructions of dSCF^BTB-I^ and dSCF^BTB-II^ suffered from a lower resolution (4.4 Å) (Figure 6C), local refinement improved the density map and enabled us to resolve the secondary structures of the two BACH1 BTB domains in the middle of the dimeric SCF (Figure 6D). In dSCF^BTB-I^, FBXL17 is engaged with BACH1-BTB through the same interface as found in mSCF^BTB^. However, in addition to the ꞵ1 strand, the α6 helix in each BACH1-BTB protomer is also structurally disordered. Remarkably, the ꞵ5 strands of the two protomers join each other and form an intermolecular anti-parallel ꞵ sheet, structurally mimicking the ꞵ1-ꞵ5 cross-protomer two-stranded ꞵ sheet found in the isolated BACH1-BTB dimer (Figure 6E). This new dimer interface is further substantiated by the cross-promoter packing of a coiled region transformed from the mostly melt α5 helices. A similar dimer interface, albeit being slightly twisted, is found in dSCF^BTB-II^. As a result, the C-terminal lobes of the two CUL1-RBX1 catalytic scaffolds become separate from each other (Figure 6C). The dimeric arrangement of these dSCF^BTB^ complexes, therefore, is predominantly, if not solely, stabilized by the BACH1 BTB domains. Surprisingly, a close inspection of the 2D class averages of the same sample revealed yet another minor population of dimeric SCF^FBXL17-BACH1-BTB^ particles, which appeared to be configured in an antiparallel manner (Figure 6A). Taken together, we postulate that these dimeric SCF^FBXL17-BACH1-BTB^ super-assemblies represent transient intermediate states of BACH1-BTB dimer remodeling, which is simultaneously catalyzed by two SCF^FBXL17^ E3 complexes.

To assess whether efficient substrate remodeling indeed requires the synergy of two copies of the E3 complexes, we isolated a BACH1-BACH2 BTB domain heterodimer with tandem affinity purification upon co-expression. Lacking the key residues for FBXL17 binding^45^, homodimeric BACH2-BTB is completely resistant to remodeling by the F-box protein (Figure 6F). Interestingly, even with an incubation time of about two days, FBXL17 can barely capture BACH1-BTB from the BACH1-BACH2 heterodimer, which is in stark contrast to the homodimeric BACH1-BTB (Figure 6G). Of note, because the BACH1-BACH2 heterodimer retains a functional ꞵ1-α6 quaternary structure degron (Figure 2F), it can still be recognized by FBXO22 (Figure 6H). These results strongly suggest that the homodimeric BACH1 BTB dimer is indeed inspected by a pair of SCF^FBXL17^, which is distinct from SCF^FBXO22^. If the structural integrity of the substrate dimer is compromised by oxidative modifications, such as S-nitrosylation, SCF^FBXL17^ can efficiently seize monomeric BACH1.

## DISCUSSION

Ubiquitin ligases have been well known to recruit substrates by recognizing their SLiM degrons^52^. Recent studies have shown that a number of substrate proteins also present their degrons via their tertiary folds^53,54^. The current study expands the degron space into the quaternary structures of protein assemblies, which allow E3s to differentiate targets based on signature “motifs” encrypted only in high-order assemblies. In the case of the ꞵ1-α6 degron of the dimeric BACH1-BTB recognized by FBXO22, substrate specificity is dictated by the proper association of the two identical subunits of the substrate. By contrast, the degradation signal of the BACH1-BTB dimer decoded by FBXL17 is determined by the strength of protein-protein interaction within the dimeric assembly, a property that can be affected by post-translation modifications associated with oxidative stress. These examples highlight the diverse nature of degrons recognized by ubiquitin ligases and underscore the complex mechanisms governing E3-substrate specificity in protein degradation. Interestingly, FBXO22 has been suggested to ubiquitinate several substrate proteins with no structural similarity to BACH1^55^. How the F-box protein recognizes these targets, presumably also through its FIST domain, remains to be determined. With a role in protein dimerization quality control, FBXL17 has been implicated in ubiquitinating and degrading a subset of BTB-domain-containing proteins when they form mismatched hetero-dimers through their BTB domains^45,46^. We propose that efficient clearance of these substrate proteins also requires synergistic actions of a pair of SCF^FBXL17^ and that the nature of the mismatched hetero-dimers recognized by FBXL17 lies in the strength of interactions between the two BTB domains. Our work indicates that a pair of SCF^FBXL17^ complexes can remodel homodimeric BACH1 when the interaction of the protomers is compromised by oxidative stress. It is conceivable that FBXL17 also surveys hetero-dimeric BTB domain-containing proteins. By probing the stability of the dimer interface, FBXL17 is able to selectively promote the degradation of either structurally compromised homodimeric BTBs or weakly associated mismatched heterodimers.

As a key transcription factor regulating heme metabolism and oxidative stress, BACH1 is tightly regulated at its protein level. FBXO22 and FBXL17 appear to have been co-evolved to play complementary roles in promoting BACH1 ubiquitination. Our studies not only reveal the distinct structural basis of BACH1 recognition by the two F-box proteins, but also shed light on the regulatory mechanisms by which different forms of oxidative stress make the transcriptional repressor discernable to the E3s (Figure 7). With an unusually large number of cysteine residues, BACH1 appears to function as a redox sensor, which can modulate its own stability in response to pro-oxidants. While heme or pro-oxidants releases BACH1 from the chromatin and exposes its ꞵ1-α6 degron to FBXO22, nitric oxide modifies at least two cysteine residues in the BACH1 BTB domain and makes the BTB dimer susceptible to FBXL17-mediated remodeling by weakening the dimer interface. Outside the BTB domain, a series of reactive cysteine residues of BACH1, including several in the bZIP domain, has been identified by proteome-wide analyses to be sensitive to modifications by H_2_O_2_ and NO donors^51,56^. It is plausible that these cysteine residues of BACH1 could also serve as sensors for a variety of pro-oxidants and possibly affect the function and stability of the transcriptional repressor via unknown mechanisms^57,58^.

**Figure 7.**
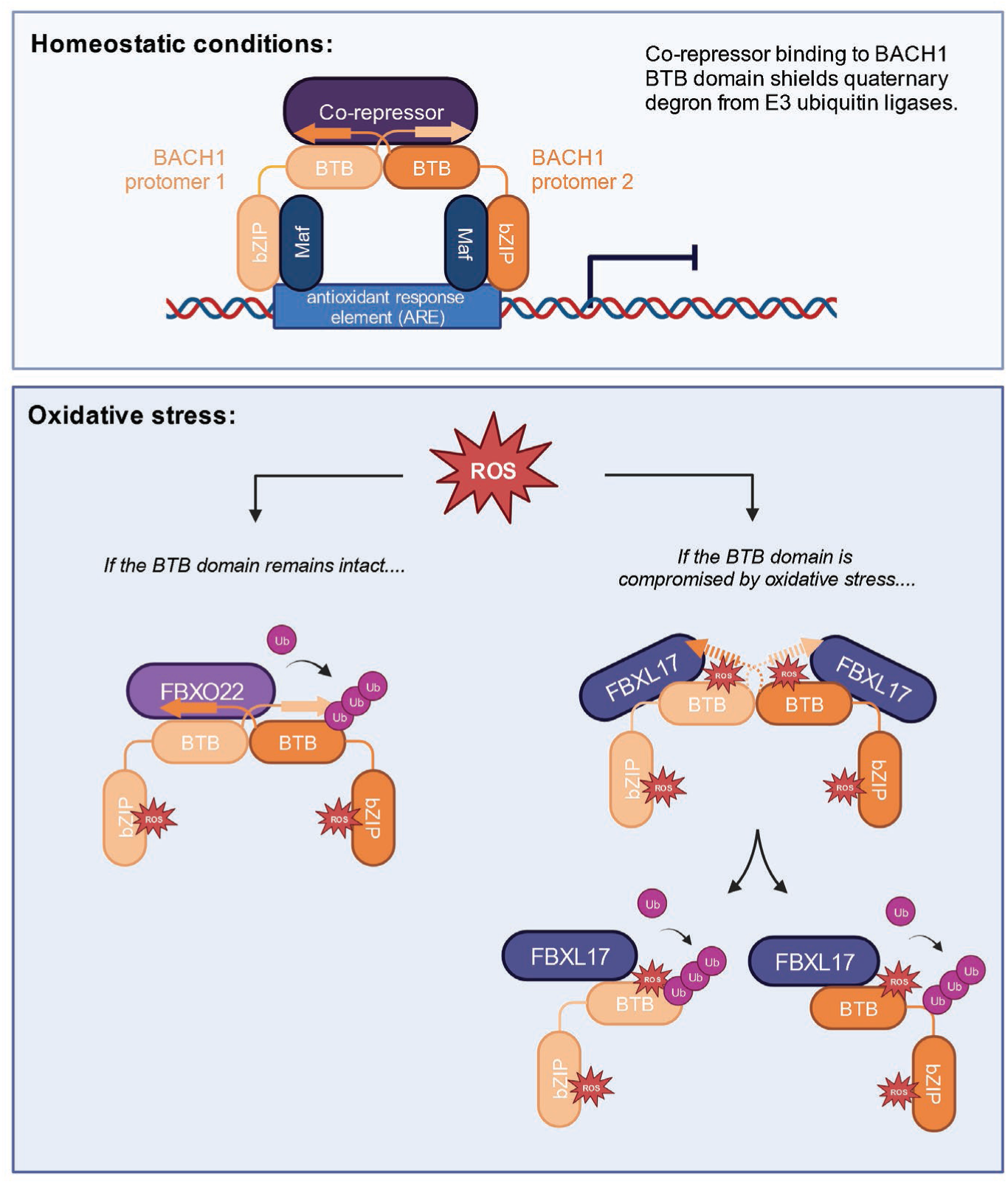
A schematic model of oxidative stress-induced BACH1 ubiquitination by FBXO22 and FBXL17. Under homeostatic conditions, BACH1 homodimerizes via its BTB domain and interacts with small MAF proteins through its bZIP domain to repress gene expression by binding to the antioxidant response elements (AREs) of oxidative stress response genes. The interaction between BACH1 and a co-repressor blocks the recognition of the ꞵ1-α6 quaternary structure degron of BACH1 by FBXO22. Under oxidative stress, reactive oxygen species (ROS) covalently modify the bZIP domain of BACH1 and release it from chromatin. If the BTB domain of BACH1 remains intact, its ꞵ1-α6 degron is recognized by FBXO22, which promotes its ubiquitination and degradation. If the pro-oxidant also modifies the BACH1 BTB domain and compromises the structural integrity of the cross-protomer ꞵ1-α6 degron, thereby, weakening the BTB dimer, a pair of FBXL17 will transiently associate with the BACH1 BTB dimer and remodel it into stably bound monomer for ubiquitination and degradation.

As a transcription factor, BACH1 has a well-established role in promoting cancer invasiveness^25-27^. In non-small lung cancer, for example, mutations in *KEAP1* and *NFE2L2* result in the stabilization of NRF2, which induces the expression of HO-1. The reduced concentration of free heme subsequently stabilizes BACH1, which in turn promotes lung cancer metastasis^20^. Our studies suggest that compounds that could modify the BTB or DNA-binding domains of BACH1 have the potential to promote the ubiquitination and degradation of BACH1 by FBXL17 or FBXO22, thereby, ameliorating BACH1-mediated cancer metastasis.

## ACKNOWLEDGMENTS

The authors would like to thank R. Yan, X. Zhao, J. Jung, and Z. Yu at the Cryo-EM Facility on the Janelia Research Campus of the Howard Hughes Medical Institute, and J.D. Quispe and S. Dickinson at the Arnold and Mabel Beckman Cryo-EM Center at the University of Washington for their assistance in electron microscopy data acquisition. The authors thank Mengxi Liu and Jorge Rivera for their contribution to this work. We also thank the NYU Proteomics Lab (supported in part by NYU School of Medicine and the Laura and Isaac Perlmutter Cancer Center Support grant P30CA016087 from the National Cancer Institute. In addition, the mass spectrometric experiments were supported with a shared instrumentation grant from the NIH, 1S10OD010582-01A1 for the purchase of an Orbitrap Eclipse) for S-nitrosylation site-mapping experiments. The authors also thank T.R. Hinds for help on BLI data analysis, H. Mao for protein expression and purification, and members of the Zheng Lab and Pagano Lab for discussion. Both N.Z. and M.P. are Howard Hughes Medical Institute Investigators. This work is also supported by NIH grant GM136250 to M.P.

## AUTHOR CONTRIBUTIONS

S.C., S.F.G, L.L., M.P., and N.Z. conceived the project. S.C. with inputs from S.F.G. constructed and purified all proteins samples in this study. S.C. performed BLI, AlphaScreen, in vitro pull-down experiments and prepared the sample for cryo-EM analysis. S.F.G. and L.L. generated CRISIPR KO cell lines, designed plasmid constructs, and performed all of the cell-based co-immunoprecipitations and protein stability assays, with notable contributions from H.V.G. and Y.K. J.P. and B.U. performed the mass spectrometric sample preparation, data acquisition and analysis. Huigang S. performed the majority of cryo-EM grid preparation, specimen screening, data collection and processing tasks with early contributions from Hui S. and S.C. The manuscript was written with inputs from all authors.

## METHODS

### Protein preparation

Human FBXO22 (N-terminally GST-tagged) and SKP1 (N-terminally His-tagged) were co-expressed and purified from Hi5 insect cells using glutathione affinity chromatography followed by TEV cleavage to remove the tags if necessary. The tag-free FBXO22-SKP1 complex was further purified by anion exchange and size exclusion chromatography. The wild-type and mutant BACH1-BTB domain (amino acids 7-128) or BACH2-BTB domain (amino acids 9-131) constructs were expressed with an N-terminal His- or GST-tag in *E. coli* BL21. The full-length BACH1 protein was constructed with an N-terminal His-Venus tag and expressed in Hi5 insect cells. His or GST-tagged BACH1 or BACH2 was purified by Ni-NTA or glutathione resin with subsequent TEV protease treatment to remove the tags if necessary. BACH1 or BACH2 was further purified by anion exchange and/or size exclusion chromatography. To immobilize BACH1-BTB for the BioLayer Interferometry (BLI) experiment, an additional Avi-tag was designed after the TEV cleavage site. After TEV cleavage of the GST-tag, Avi-BACH1-BTB was generated and subsequently biotinylated in a biotin ligase BirA-catalyzed biotinylation reaction. Biotin-BACH1-BTB was further purified by size exclusion chromatography. GLMN with an N-terminal GST tag was expressed and purified from *E. coli* BL21 using glutathione affinity chromatography followed by TEV cleavage to remove the tag. Tag-free GLMN was further purified by anion exchange and size exclusion chromatography. RBX1 (amino acids 16-108) and CUL1 with two short unstructured segments (amino acids 1-12 and 58-81) truncated were constructed with an N-terminal His-tag followed by a TEV cleavage site. After co-expression in *E. coli* BL21(DE3), the CUL1-RBX1 complex was purified by a Ni-NTA affinity column followed by TEV cleavage to remove the tag. The tag-free CUL1-RBX1 was further purified by gel filtration. Separately purified BACH1-BTB, FBXO22-SKP1, CUL1-RBX1, and GLMN were mixed at the molar ratio of 3:1:1:1.5 with BACH1 and GLMN at extra to assemble and purify the GLMN-SCF^FBXO22-BACH1^ complex on size exclusion chromatography for cryo-EM analysis. FBXL17 (amino acids 310–701) with an N-terminal His-MBP-tag and SKP1 with an N-terminal His-tag were co-expressed and purified from Hi5 insect cells using Ni-NTA affinity chromatography followed by TEV cleavage to remove the tags if necessary. GST-BACH1-BTB without or with NOR3-S1PC treatment and FBXL17-SKP1 were incubated at various times and purified by a glutathione affinity column. Thereafter, CUL1-RBX1 and GLMN were loaded on the above column to form the GLMN-SCF^FXL17-BACH1-BTB^ complex. After on-column cleavage by TEV, the tag-free complex was collected in the flow-through of the glutathione affinity column and was further purified by size exclusion chromatography for cryo-EM analysis.

### Affinity pull-down assay

The GST and Ni^2+^ pull-down assays were performed using ∼50 μg of purified GST-or His-tagged protein as the bait. Their binding partners were applied in excess. BACH1-BTB was pre-treated with the saturating amount of NOR3 and S1PC at 100 µM overnight at 4°C if indicated. Reaction mixtures with various incubation times were incubated with 20 μL glutathione or Ni^2+^ beads at 4°C with gentle shaking. After extensive wash with the buffer containing 20 mM Tris, pH 8.0, 150 mM NaCl, 0.25 mM TCEP (additional 20 mM imidazole for the Ni^2+^ pull-down assay), the protein complexes on the beads were eluted by 40 µl 20 mM glutathione or 300 mM imidazole. The eluted samples were resolved by SDS-PAGE and analyzed by Coomassie staining.

### BioLayer interferometry

The binding affinity between FBXO22 (or FBXL17) and BACH1-BTB was measured using the Octet Red 96 (ForteBio, Pall Life Sciences) following the manufacturer’s procedures. The optical probes were coated with streptavidin, loaded with 200 nM biotinylated BACH1-BTB (without or with NOR3-S1PC treatment), or anti-GST probes loaded with 200 nM GST-FBXO22-SKP1. Subsequently, the probes were quenched with 0.5 mM biocytin or 1 µM GST protein prior to kinetic binding analysis. The reactions were carried out in black 96 well plates maintained at 30°C. The reaction volume was 200 μL in each well. The binding buffer contained 25 mM HEPES, pH 7.4, 100 mM NaCl, 1 mM TCEP, 0.1% Tween-20, and 0.05 mg/mL Bovine Serum Albumin. As the analyte, BACH1-BTB or FBXL17-SKP1 was tested at various concentrations. There was no binding of the analyte to the unloaded probes.

Binding kinetics of the analyte at different concentrations were measured simultaneously. The data were analyzed by the Octet data analysis software. The dissociation constant (K_d_) was determined from the steady-state equilibrium measurements. All BLI experiments have been repeated at least 2-3 times.

### Amplified luminescence proximity homogeneous assay

AlphaScreen assays for measuring protein-protein interactions were performed using EnSpire reader (PerkinElmer). Biotinylated BACH1-BTB was immobilized to streptavidin-coated AlphaScreen donor beads. His-SKP1-FBXO22 was bound to anti-His acceptor beads. The acceptor and donor beads were brought into proximity by the interactions between the E3 and its substrate. Excitation of the donor beads by a laser beam of 680 nm promotes the formation of singlet oxygen. When an acceptor bead is in close proximity, the singlet oxygen reacts with thioxene derivatives in the acceptor beads and causes the emission of 520-620 nm photons, which are detected as the binding signal. If the beads are not in close proximity to each other, the oxygen will return to its ground state and the acceptor beads will not emit light. Competition assays were performed by titrating GST-tagged NCOR1 fragments or GST alone at various concentrations. To establish the binding signal, the concentration of tagged E3 and biotinylated BACH1-BTB were first titrated with fixed amount of donor and acceptor beads. The concentration of each component was selected based on the rising AlphaScreen signal before the beads became saturated. 25 nM His-SKP1-FBXO22 and 8.3 nM biotinylated BACH1-BTB were used. The binding assay buffer contained 25 mM HEPES, pH 7.4, 100 mM NaCl, 1 mM TCEP, 0.1% Tween-20, and 0.05 mg/mL Bovine Serum Albumin. 5 μg/mL donor and acceptor beads were used in the assays. The experiments were performed in three or four replicates. IC_50_ was determined using non-linear curve fitting of the dose response curves generated with Prism 7 (GraphPad).

### SEC–MALS

Size-exclusion chromatography coupled with light scattering, refractive index, and ultraviolet absorption (SEC-LS-RI-UV) was done using a SEC-MALS system consisting of an Agilent HPLC pump with a 1260 Infinity lamp, a Dawn light scattering instrument (Wyatt), an OptiLab refractive index instrument (Wyatt), and the Superdex 200 10/300 GL (GE Healthcare). The column was equilibrated with 20 mM Tris-HCl pH 8.0, 150 mM NaCl, and 0.25 mM TCEP. Tag-free FBXO22-SKP1, His-BACH1-BTB and their mixture were analyzed at room temperature. Similarly, His-MBP-FBXL17-SKP1, NOR3-S1PC-treated His-BACH1-BTB and their mixture with different incubation times were analyzed.

### In vitro ubiquitination assay

A reaction mixture containing 4 μM His-Venus-BACH1 (full length), 3 μM FBXO22-SKP1, 1 μM NEDD8∼CUL1-RBX1, 1 μM UBCH5 and CDC34, 0.4 μM UBE1 and 70 μM ubiquitin and 2 mM ATP/10 mM MgCl_2_ was incubated at 37 °C for 1.5 h with multiple controls, each lacking one component of the reaction. The reaction mixtures were resolved by a 10% SDS–PAGE gel. The BACH1 signal were monitored by detecting the Venus-fluorescence absorbance at 488 nm. The absorbance at 647 nm was used to detect the protein markers.

### Cryo-EM sample preparation and data collection

The GLMN-SCF^FBXO22-BACH1-BTB^ complex and the GLMN-SCF^FXL17-BACH1-BTB^ complex (without or with NOR3-S1PC treatment with different incubation times) were concentrated to ∼2.0 mg/ml for cryo-EM sample preparation. For each complex, ∼3 μL sample was applied to a glow-discharged Quantifoil R1.2/1.3 200 mesh copper grid. The grid was subsequently blotted for ∼3 s (blotting force at -2, temperature at 10 °C and relative humidity at 100%), plunged and flash frozen into liquid ethane using a Vitrobot Mark IV system (Thermo Fisher Scientific), and stored in liquid nitrogen for data collection. The grids were inspected and screened with Talos Glacios equipped with a K3 camera. For the GLMN-SCF^FBXO22-BACH1-BTB^ sample, data collection was carried out on a Titan Krios transmission electron microscope (Thermo Fisher Scientific) operated at 300 kV at the University of Washington. The automation scheme was implemented using the Leginon software^61^ at a nominal magnification of 105 K, resulting in a physical pixel size of 0.84 Å. The zero-loss-energy images were acquired on a Gatan K3 direct detector operated in super-resolution counting mode (pixel size: 0.42 Å) with the slit width of post-column Gatan BioQuantum GIF energy filter set to be 20 eV. The dose rate was adjusted to 13.9 e^-^/Å^2^S, and a total dose of 59 e^-^/Å^2^ for each image fractionated into 50 frames. The images were recorded at a defocus range of -0.8∼3.5 μm. For the chemical-treated or untreated GLMN-SCF^FXL17-BACH1-BTB^ complex, data collection was carried out on a 300 kV FEI Titan Krios transmission electron microscope operated at 300 kV at the HHMI Janelia Research Campus. The automated data collection was carried out with SerialEM software^62^ using beam-image shift^63^ at a nominal magnification of 165 K. The images were acquired on Falcon 4i direct electron detector operated in electron counting mode (pixel size: 0.743 Å) with the slit width of Selectris X (Thermo Fisher Scientific) energy filter set to be 6 eV. The dose rate was set to 15.39 e^-^/Å^2^S, and a total dose of 60 e^-^/Å^2^ for each image fractionated into EER (Electron-event representation) frames^64^. The images were recorded at a defocus range of 0.8∼1.5 μm.

### Image processing and 3D reconstruction

For the GLMN-SCF^FBXO22-BACH1-BTB^ sample, 6,131 movies were acquired. The beam-induced motion of each micrograph stack was corrected by MotionCor2^65^. The motion-corrected micrographs were imported into CryoSPARC for the following processing^66^. The defocus parameter of each motion-corrected micrograph was determined by Gctf^67^. 4,991 micrographs were kept after filtering the micrographs with CTF parameters and visual inspection. An *ab-initio* reconstruction from a previous data collection was used to generate 2D projections as templates for particle picking. 2,888,035 particles were picked, extracted, and subjected to 2D classification. After two rounds of 2D classification, 243,712 particles were selected as training particles for Topaz particle picking^68^. 3,645,675 particles were picked with Topaz. After data cleaning by 2D classification, 949,569 particles were kept and subjected to *ab-initio* reconstruction. Subsequently, all the particles were used for heterogenous refinement. 413,179 particles from good reconstruction were selected for non-uniform refinement^69^ to generate a reconstruction with an overall resolution of 3.9 Å. To better resolve the interface between the substrate and its receptor, a soft mask focused on BACH1-BTB and FBXO22-SKP1 was applied to 3D Variability Analysis^70^. 171,378 particles were chosen for local refinement. The BACH1-BTB-FBXO22-SKP1 subcomplex was refined to 3.8 Å. More details about the data processing can be found in Figure. S1.

For the native GLMN-SCF^FXL17-BACH1-BTB^ sample, 12,618 movies were collected. The beam-induced motion of micrographs was corrected by Patch Motion Correction in CryoSPRAC. The defocus of motion-corrected micrographs was estimated by CTFFIND4^71^. 12,423 micrographs were kept while the rest were rejected by CTF parameters. These micrographs were subjected to Blob particle picker and two rounds of 2D classification sequentially. 497,102 out of 3,087,124 particles were picked for heterogenous refinement using reconstructions from the previous dataset as templates. 267,523 good particles from the refinement were selected and further cleaned by 2D classification. 84,154 particles were chosen as training particles for Topaz particle picking. 7,309,475 particles were picked with Topaz and subjected to two rounds of 2D classification. 1,099,238 particles were selected for *ab-initio* reconstruction and heterogeneous refinement sequentially. 364,786 particles from one good class were applied to non-uniform refinement. The overall resolution of the complex in the monomeric form was refined to 3.1 Å. To improve the local density of FBXL17-BACH1-BTB, all particles were subjected to 3D Variability. 193,908 particles were selected for local refinement to generate the SKP1-FBXL17-BACH1-BTB subcomplex with a resolution of 3.2 Å. In addition to the above complex in the monomeric form, the dimeric GLMN-SCF^FXL17-BACH1-BTB^ complex was detected as well. 30,123 particles of the dimer were selected as training particles for Topaz particle picking. 244,189 out of 2,969,728 particles were chosen for heterogenous refinement. Two good classes representing different conformations, named dSCF^BTB-I^ and dSCF^BTB-II^, were kept with 54,546 particles and 47,336 particles, respectively. The C2 symmetry was imposed for non-uniform refinement. The overall resolution of the two classes reached 4.4 Å and 4.5 Å, respectively. To overcome the pseudo-symmetry of the dimeric reconstructions, all particles of dSCF^BTB-I^ were subjected to symmetry expansion in CryoSPARC. Afterward, a soft mask focused on the leucine-rich repeat domain of FBXL17 and BACH1-BTB was applied to subtract the signal from raw particles. These subtracted particles were used for local refinement. The density of the FBXL17-LRR-BACH1-BTB structure was refined to 4.1 Å. See Figure S5 for more details.

For the compound-treated GLMN-SCF^FXL17-BACH1-BTB^ sample, the collected data was processed with CryoSPARC in a similar manner. The data processing workflow was shown in Figure S4. Briefly, 22,720 EER movies were collected and subjected to patch motion correction and patch CTF estimation sequentially. After filtering with CTF parameters, 20,028 micrographs were kept for the subsequent blob picker and 2D classification.

Representative particles were selected as the training data for Topaz particle picking. After heterogeneous refinement, a monomeric SCF^FXL17-BACH1^ class yielded a 3D reconstruction at 5.7 Å resolution. A dimeric SCF^FXL17-BACH1^ with 461 particles was selected and employed for Topaz particle picking. After 2D classification and heterogeneous refinement, two good classes of dimeric SCF^FXL17-BACH1^ were kept. 10,173 particles and 7,944 particles from dimeric reconstructions were subjected to non-uniform refinement with C2 symmetry imposed. After refinement, the overall resolution of two classes both reached 7.8 Å.

### Model building and refinement

The initial structural model of FBXO22-SKP1-BACH1-BTB was predicted with AlphaFold-Multimer in Google ColabFold^72^ and fitted into the 3.8 Å Cryo-EM map by using UCSF Chimera^73^. Subsequently, the model was inspected and trimmed in Coot^74^ based on the protein sequences and the EM density. The model was further improved by real-space refinement in PHENIX^75,76^ and manual rebuilding in Coot. To generate an initial model of the monomeric SCF^FBXL17-BACH1-BTB^ complex, BACH1-BTB (PDB: 2IHC) was applied to superimpose with and replace the KEAP1-BTB subunit of the SCF^FBXL17-KEAP1-BTB^ complex (PDB: 6WCQ) in PyMOL. This model was fitted into the 3.2 Å cryo-EM map using UCSF Chimera. The BACH1 BTB domain was manually adjusted to fit the density in Coot. The model was further improved by using real-space refinement in PHENIX and manual rebuilding in Coot. For the SCF^FBXL17-BACH1^ dimer, two copies of the monomeric complexes were fitted into the 4.4 Å cryo-EM maps and subsequently trimmed and adjusted in Coot. The dimeric interface of BACH1-BTB in the SCF complex was rebuilt in Coot since it is completely different from that of the isolated BACH1-BTB dimer. The model of dimeric FBXL17-BACH1-BTB was refined in PHENIX by real-space refinement. Cryo-EM data collection and refinement statistics for all structures in this paper were summarized in Table S1.

### Plasmids and antibodies

BACH1, FBXO22, and FBXL17 cDNAs were inserted into a modified pcDNA3.1 vectors containing a variety of N-terminal tags, including 2xFLAG tag-2x Strep tag, and HA tag via sub-cloning. Site-directed mutagenesis was performed using Q5 High-Fidelity DNA polymerase (New England Biolabs (NEB)) with primers generated using either QuikChange Primer Design (Agilent) or NEBaseChanger (New England Biolabs (NEB)). The following antibodies were used in this study: vinculin (1:1000, Bethyl), Bach1 (1:1000, Bethyl), HO-1 (1:1000, Bethyl), FBXO22 (1:1000, Proteintech), Nrf2 (1:1000, Cell Signaling), Keap1 (1:10 000, Proteintech), HA (1:1000, Sigma-Alrdich), FLAG (1:4000, Sigma-Aldrich), Skp1 (1:5000, Michele Pagano’s lab), NCoR1 (1:1000, Cell Signaling Technology), MafF (1:1000, Proteintech), PCNA (1:2000, Santa Cruz Biotechnology).

### Cell Culture

Cell lines were purchased from ATCC and routinely monitored for *Mycoplasma* contamination using the Universal Mycoplasma Detection Kit (ATCC 30-1012K). HCT-116 (ATCC CCL-247) cells were propagated in McCoy’s 5A medium (Gibco). HEK293T (ATCC CRL-3216) cells and FLAG-FBXL17 knock-in HEK293T cells (provided by Michael Rape lab) were propagated in Dulbecco’s modified Eagle’s medium (DMEM) (Gibco). Tissue culture media were supplemented with 10% fetal bovine serum (FBS) (Corning Life Sciences) and 1% penicillin-streptomycin (Corning Life Sciences). Cells were maintained at 37^°^C and 5% CO_2_ in a humidified atmosphere. MLN-4924 (Active Biochem) was used at 2 μM. MG-132 (Peptides International) was used at 10 μM. Spermine NONOate (sNONO) (Cayman Biochem) was prepared in 0.01N NaOH and used at 100 μM. S-nitrosoglutathione (GSNO) (Sigma-Aldrich) was prepared in DMSO and used at 100 μM. NOR3 (Sigma-Aldrich) and S-1-propenyl-L-cysteine (S1PC) (MedChemExpress) were prepared in DMSO immediately prior to the experiment and used at 25 μM (NOR3) and 50 μM (S1PC) respectively.

### Co-immunoprecipitation assay

HEK293T cells were transiently transfected using polyethylenimine (Polysciences); HCT-116 cells were transiently transfected with Lipofectamine 3000 (Thermo Fisher Scientific). Where indicated, 48 h after transfection, cells were treated with 2.5 μM MLN-4924 for 3 h before further treatment with either vehicle (DMSO) or the indiciated NO donor. Cells were collected by scraping and washed with PBS prior to lysis in IP lysis buffer (50 mM Tris-HCl, pH 7.5, 150 mM NaCl, 1 mM EDTA, 1 mM EGTA, 5 mM MgCl_2_, 0.2% NP-40) supplemented with protease inhibitor (Complete ULTRA, Roche) and phosphatase inhibitor (Phosphatase Inhibitor Cocktail 2, Sigma-Aldrich) immediately before the experiment. The supernatant was clarified by centrifugation at 21, 000 x g for 10 minutes at 4^°^C. For FLAG-tagged proteins, the supernatant was incubated with FLAG-M2 agarose beads (Sigma-Aldrich) for 2h at 4^°^C. For HA-tagged proteins, the supernatant was incubated with Pierce Anti-HA magnetic beads (Thermo Fisher Scientific) for 2h at 4^°^C. The beads were then washed 5 times in lysis buffer before incubating the beads in NuPAGE LDS Sample Buffer (Thermo Fisher Scientific) supplemented with ꞵ-mercaptoethanol (Sigma-Aldrich) at 95^°^C for 10 minutes. For Western blotting, samples of the whole cell extract and immunoprecipitates were separated by SDS-PAGE and transferred onto 0.45 μm Immobilon-P PVDF membranes (Millipore Sigma). Membranes were then blocked in 5% nonfat dried milk / PBST for 30 minutes at room temperature and incubated with the indicated primary antibodies at 4^°^C overnight. After washing the membranes, secondary antibodies conjugated with horseradish peroxidase were applied. Immunoreactive bands were visualized via application of enhanced chemiluminescence (ECL) reagent (Thermo Fisher Scientific).

### Gene silencing by siRNA

ON-TARGETplus SMARTpool siRNA oligos targeting human *FBXL17* (Dharmacon, cat. no. L-024385-00-0005) and ON-TARGETplus non-targeting siRNA #1 (Dharmacon, cat. no. D-001810-01) as a negative control were used in this study. The siRNA oligos were transfected using Lipofectamine RNAiMAX Transfection Reagent (Thermo Fisher Scientific). To achieve satisfactory siRNA-mediated silencing, siRNA oligonucleotides were transfected for two to three rounds.

### Cell-based protein stability assay

HCT-116 cells were seeded to about 50% confluency the day before treatment. Cells were treated with 100 μM spermine NONOate (sNONO) for 6h. Cells were then washed with ice-cold PBS supplemented with 100 μM phenylmethanesulfonyl fluoride (PMSF) (Sigma-Aldrich) before they were collected by scraping. Cell pellets were lysed in RIPA buffer (Pierce) supplemented with protease inhibitor (Complete ULTRA, Roche) and phosphatase inhibitor (Phosphatase Inhibitor Cocktail 2, Sigma-Aldrich) immediately before the experiment. The supernatant was clarified by centrifugation at 21, 000 x g for 10 minutes at 4^°^C. Protein concentrations in cell lysates were then normalized with the Pierce BCA Protein Assay Kit (Thermo Fisher Scientific), according to the manufacturers’ instructions. Samples of the cell lysates were then prepared with NuPAGE LDS Sample Buffer (Thermo Fisher Scientific) supplemented with ꞵ-mercaptoethanol (Sigma-Aldrich) and changes in protein levels were visualized via Western blot.

### CRISPR-Cas9 genome editing

To generate *FBXO22* knockout cells, optimal sgRNA target sequences were designed using the Benchling CRISPR Genome Engineering tool (https://www.benchling.com). *FBXO22* gRNA (CTTCGTGTTGAGTAACCTGG) was cloned into pSpCas9(BB)-2A-GFP (px458), a gift from F. Zhang (Addgene plasmid #48138), using the following primers: CACCGCTTCGTGTTGAGTAACCTGG and aaacCCAGGTTACTCAACACGAAGC. 5x10^6^ HCT-116 cells were seeded into a 15 cm dish before being transfected with 10 μg sgRNA-containing px458 plasmid using Lipofectamine 3000 (Thermo Fisher Scientific). 48h after transfection, the top 5% of GFP-positive cells were sorted using the Beckman Coulter MoFlo XDP cell sorter (100 μm nozzle). 10 000 cells were plated between two 15 cm dishes (5000 cells per plate) and grown for about 10-14 days to allow single-cell clones to form colonies. The colonies were then picked and plated into individual wells of a 96-well plate. After 48 hours, the 96-well plate were split into two new 96-well plates, with one plate used for genotyping. For genotyping, genomic DNA was extracted using QuickExtract (Epicentre) and genotyping PCRs were performed with MyTaq HS Red Mix (Bioline) using the following *FBXO22* genotyping primers: GCAACCCTATGCTGGCGTAA and AGATACCTCGCGGGTAGGG.

The resulting PCR products were then purified and sequenced to determine the presence of a CRISPR-mediated frameshift event that results in an early stop codon. Positive knockout candidates were further validated by Western blot. Five knockout clones were then pooled together to mitigate any clonal effects, and this collection of pooled knockout cells were propagated and subsequently used for experiments.

### Biotin Switch Assay

Cells were treated with either 100 μM GSNO or 100 μM sNONO for 1h prior to collection. Cells were then subjected to the S-nitrosylation Protein Detection Kit (Cayman Chemical), according to the manufacturers’ instructions. Briefly, free reactive thiol groups were blocked with N-ethylmaleimide (NEM) for 30 minutes at room temperature, then excess NEM was removed via acetone precipitation. Any S-nitrosylated cysteines were subsequently reduced and labelled with a sulfhydryl-specific biotin. Excess reducing and labelling were removed via acetone precipitation. The final protein pellet was then resuspended in IP lysis buffer (50 mM Tris-HCl, pH 7.5, 150 mM NaCl, 1 mM EDTA, 1 mM EGTA, 5 mM MgCl_2_, 0.2% NP-40, 1 mM DTT) supplemented with protease inhibitor (Complete ULTRA, Roche) and phosphatase inhibitor (Phosphatase Inhibitor Cocktail 2, Sigma-Aldrich). The resuspended protein was incubated with streptavidin Dynabeads MyOne Streptavidin C1 beads (Thermo Fisher Scientific) for 2h at 4°C before the beads were washed 4 times with IP lysis buffer. Samples of the resuspended protein and the streptavidin bead-bound protein were prepared in NuPAGE LDS Sample Buffer (Thermo Fisher Scientific) supplemented with ꞵ-mercaptoethanol (Sigma-Aldrich) and proteins were assessed via Western blot.

### S-nitrosylation site mapping

10 μg of recombinantly purified BACH1 BTB was allowed to react in the presence of 100 μM NOR3 and 100 μM S1PC for 30 minutes at room temperature before excessing nitric oxide donor was removed by Zeba desalting column pre-equilibrated with 50 mM Tris-HCl (pH 8.0), 150 mM NaCl. Nitrosylated protein was subsequently treated with the S-nitrosylation Protein Detection Kit (Cayman Chemical), according to the manufacturers’ instructions. Biotin switch-treated protein reduced with 2.5 μL of 0.2 M dithiothreitol (Sigma) for 1 hr at 57°C and the samples cooled before adding 2.5 μL of 0.5M iodoacetamide and incubated in the dark for 45 minutes. The sample was digested with 400 ng of trypsin overnight at room temperature and light agitation. The digestion was stopped by adding 10% trifluoroacetic acid (TFA) to a final concentration of 0.5% TFA. The resulting peptides were desalted using equilibrated Pierce™C18 Spin columns. The samples were washed 3 times with 0.1% TFA followed by a wash with 0.5% acetic acid. The desalted peptides were eluted using 80% acetonitrile in 0.5% acetic acid. The peptides were dried using a SpeedVac concentrator. Samples were reconstituted in 0.5% acetic acid and stored at -80°C until further analysis. 40 pmol of each sample was loaded onto an Acclaim PepMap trap column (2 cm x 75 µm) in line with an EASY-Spray analytical column (50 cm x 75 µm ID PepMap C18, 2 μm bead size) using the autosampler of an EASY-nLC 1200 HPLC (Thermo Fisher Scientific). Solvent A consists of 2% acetonitrile in 0.5% acetic acid, and Solvent B of 80 % acetonitrile in 0.5% acetic acid. Peptides were gradient eluted with a flowrate of 200nL/min into an Orbitrap Eclipse Tribrid (Thermo Fisher Scientific) mass spectrometer. The gradient used was the following: in 5 min to 5% solvent B, 60 min to 35% solvent B, 10 min to 45% solvent B, and 10 min to 100% solvent B. High resolution full MS spectra were acquired with a resolution of 120,00, an AGC target of 4e5, with a maximum ion time of 50 ms, and a scan range of 400 to 1500 m/z. All HCD MS/MS spectra were collected with the following instrument parameters: resolution of 30,000, AGC target of 2e5, maximum ion of 200ms, one microscan, 2 m/z isolation window, and NCE of 27. The resulting MS/MS spectra were searched against BACH1 BTB and common contaminants using the search engine Byonic (Protein Metrics). Settings were as following: trypsin cleavage allowing for 2 missed cleavages, precursor, and fragment ion mass tolerance of 10 ppm, variable modifications of carbamidomethyl, N-ethylmaleimide, and 3-(N-Maleimidopropionyl)-biocytin on cysteine, deamidation on asparagine and glutamine, and oxidation on methionine. Spectra identified as containing modified cysteine residues were manually verified.

## Supplementary Materials

Cao S. *et al.* 2023

## Supplementary Figure Legends

**Figure S1.**
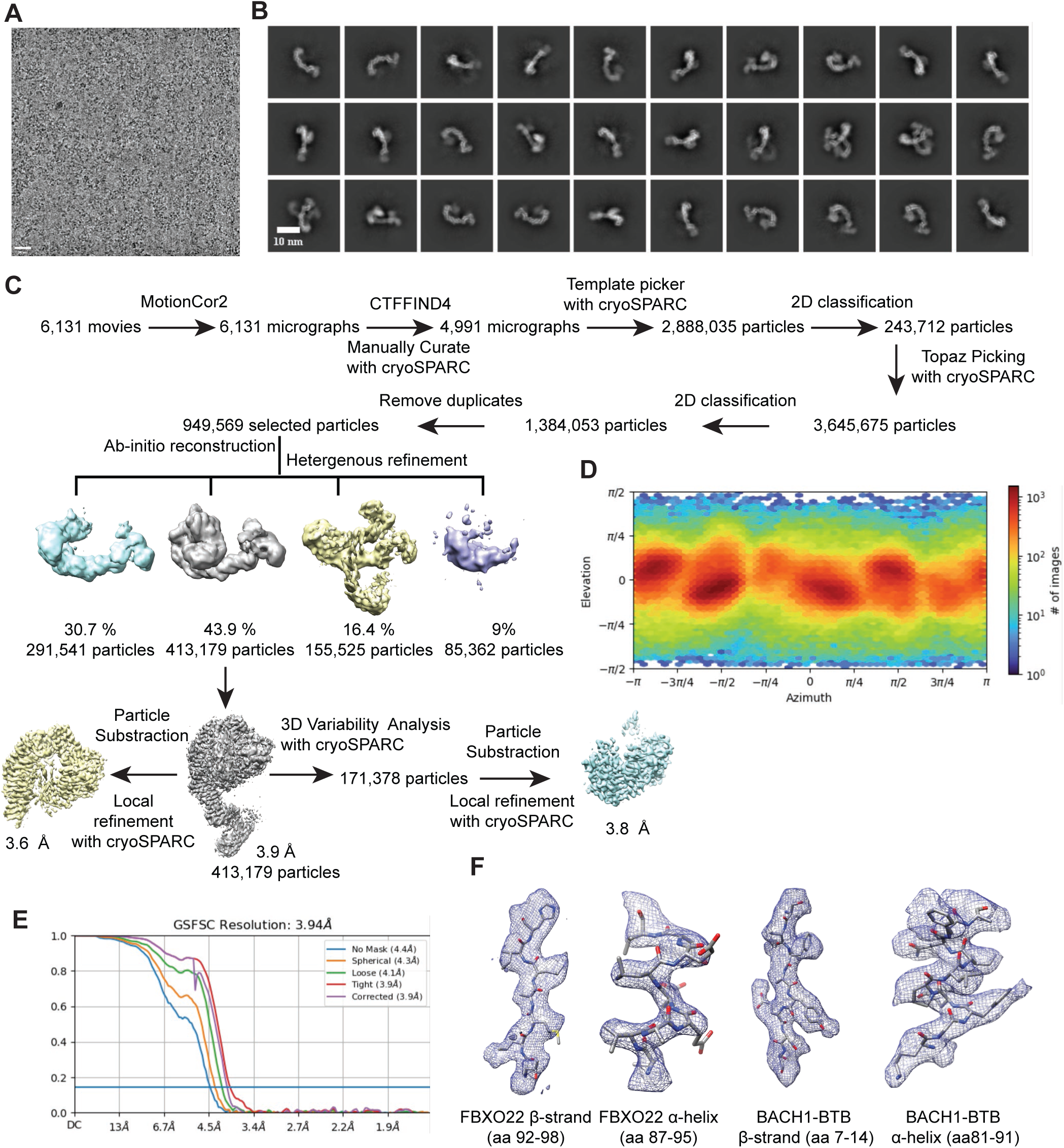
The schematic workflow of single particle reconstruction of the SCF^FBXO22-BACH1^complex. A. A representative cryo-EM micrograph. **B.** Typical 2D averages of the cryo-EM dataset. Scale bar 10 nm. **C.** The flowchart of single particle analysis of the SCF^FBXO22-BACH1^complex. **D.** The angular distribution of particles used in the final reconstruction. **E,** Fourier shell correlation (FSC) curves for SCF^FBXO22-BACH1^. At the Gold-standard threshold of 0.143, the resolution is 3.9 Å. **F.** Representative density in local refined EM map.

**Figure S2.**
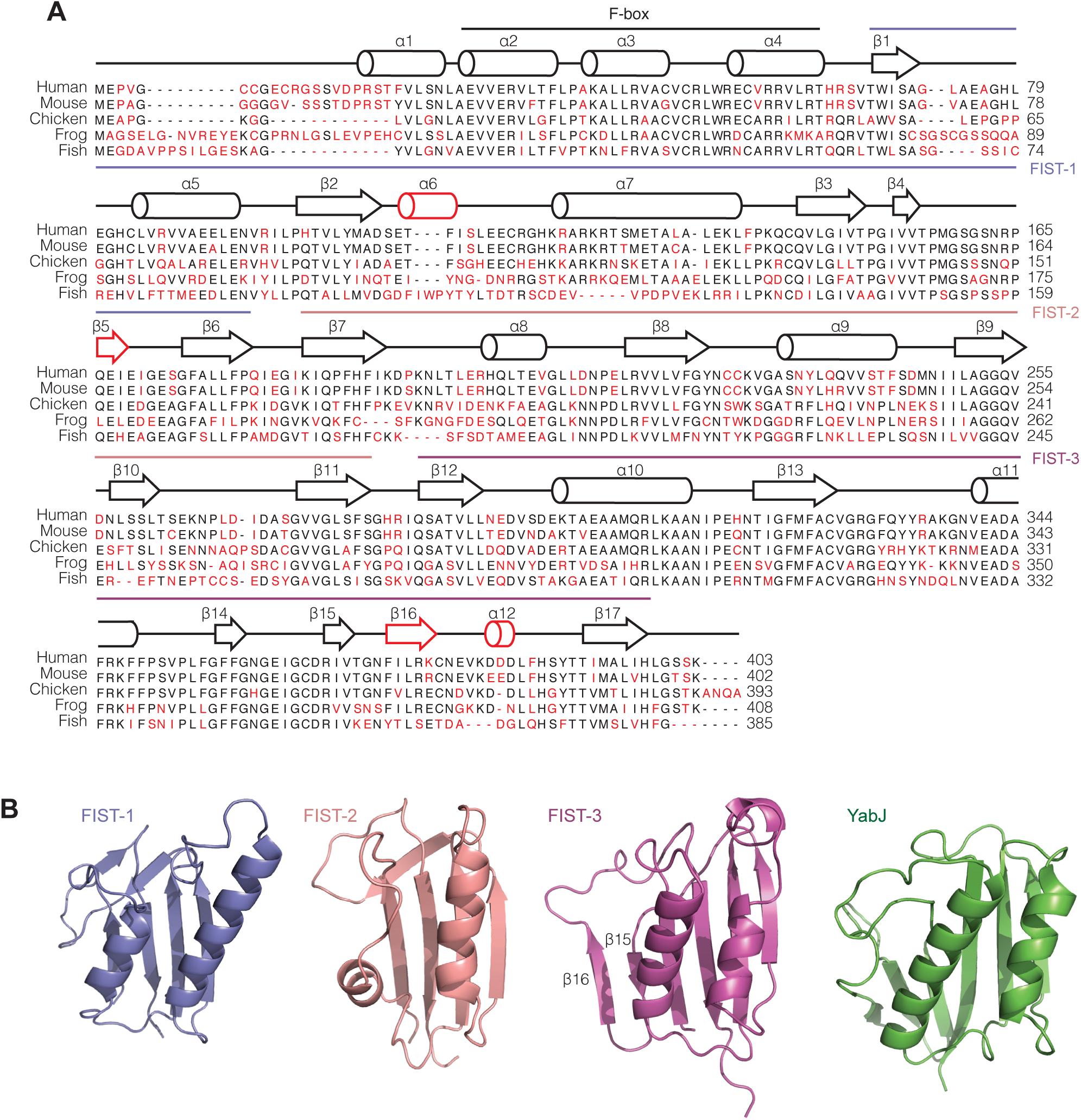
Sequence and structural analysis of FBXO22. A. Sequence alignment of five vertebrate FBXO22 orthologues with second structure annotations. The sequences of the three FIST domains are underlined in different colors (slate, salmon, and purple). Unique secondary structure elements in each repeat are highlighted in red. **B.** A comparison of the three FIST domains of FBXO22 and homologous structure of YabJ from the YjgF superfamily (PDB:1QD9).

**Figure S3.**
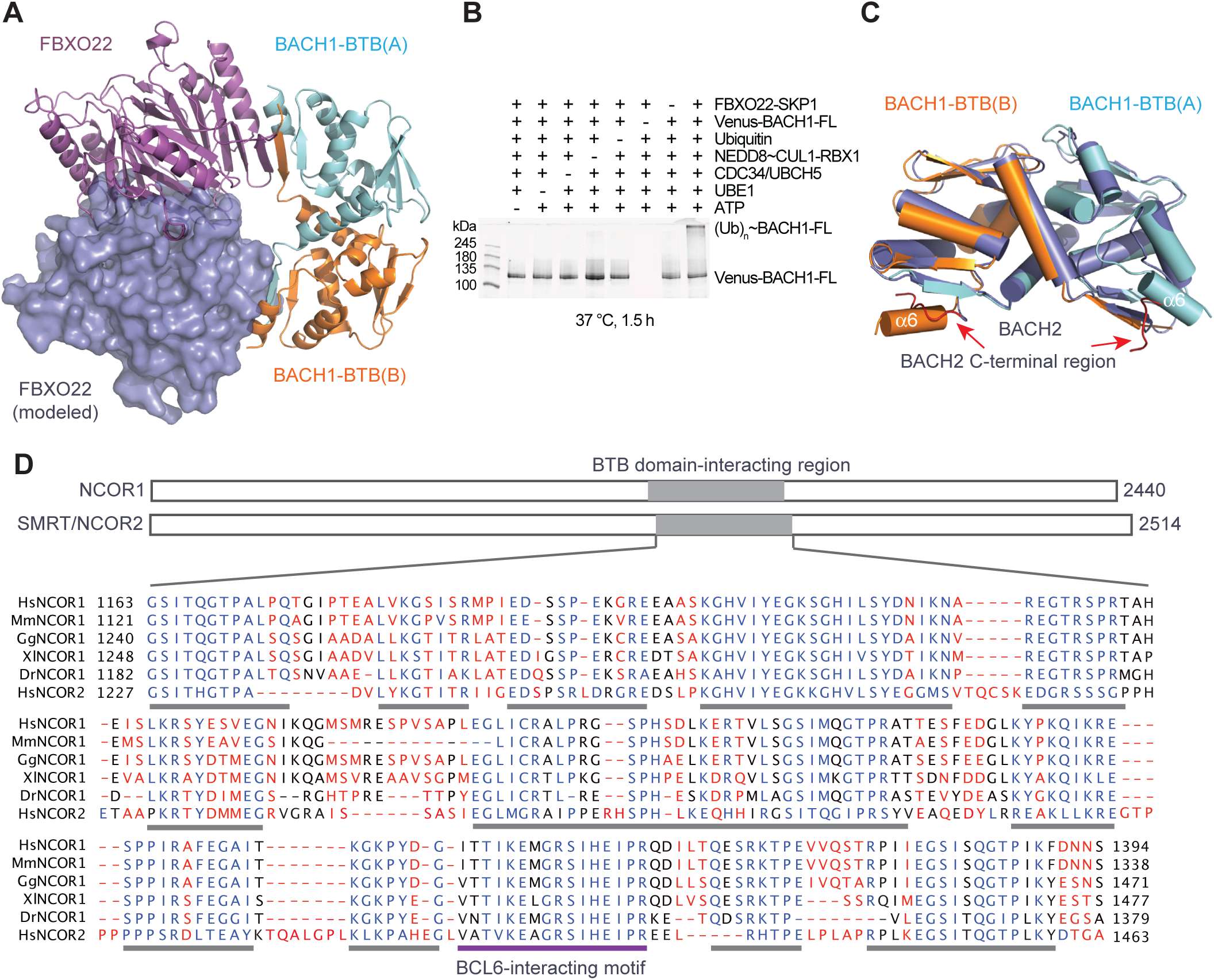
Structural and biochemical analyses of FBXO22-BACH1 interaction and sequence analysis of NCOR1/2. A. Steric hindrance prevents the formation of a BACH1-BTB-FBXO22 complex with a 2:2 ratio. The asymmetric complex formed between a BACH1 dimer (protomer A: light blue; subunit B: orange) and FBXO22 (purple) is shown as cartoon diagram. A second copy of FBXO22 shown in slate surface representation is modeled onto the BACH1-BTB dimer and is in clash with the other FBXO22 macromolecule. **B.** *In vitro* ubiquitination of BACH1 by SCF^FBXO22^. **C.** Superposition of the crystal structures of BACH1-BTB (PDB:2IHC) and BACH2-BTB (PDB:3OHU). BACH2 C-terminal region is disordered and highlighted in red. **D.** Sequence alignment of five NCOR1 vertebrate orthologs and human NCOR2. Highly conserved short linear motifs (SLiMs) are underlined with gray bars. The BCL6 BTB-interacting motif found in PDB:1R2B is underlined with a purple bar.

**Figure S4.**
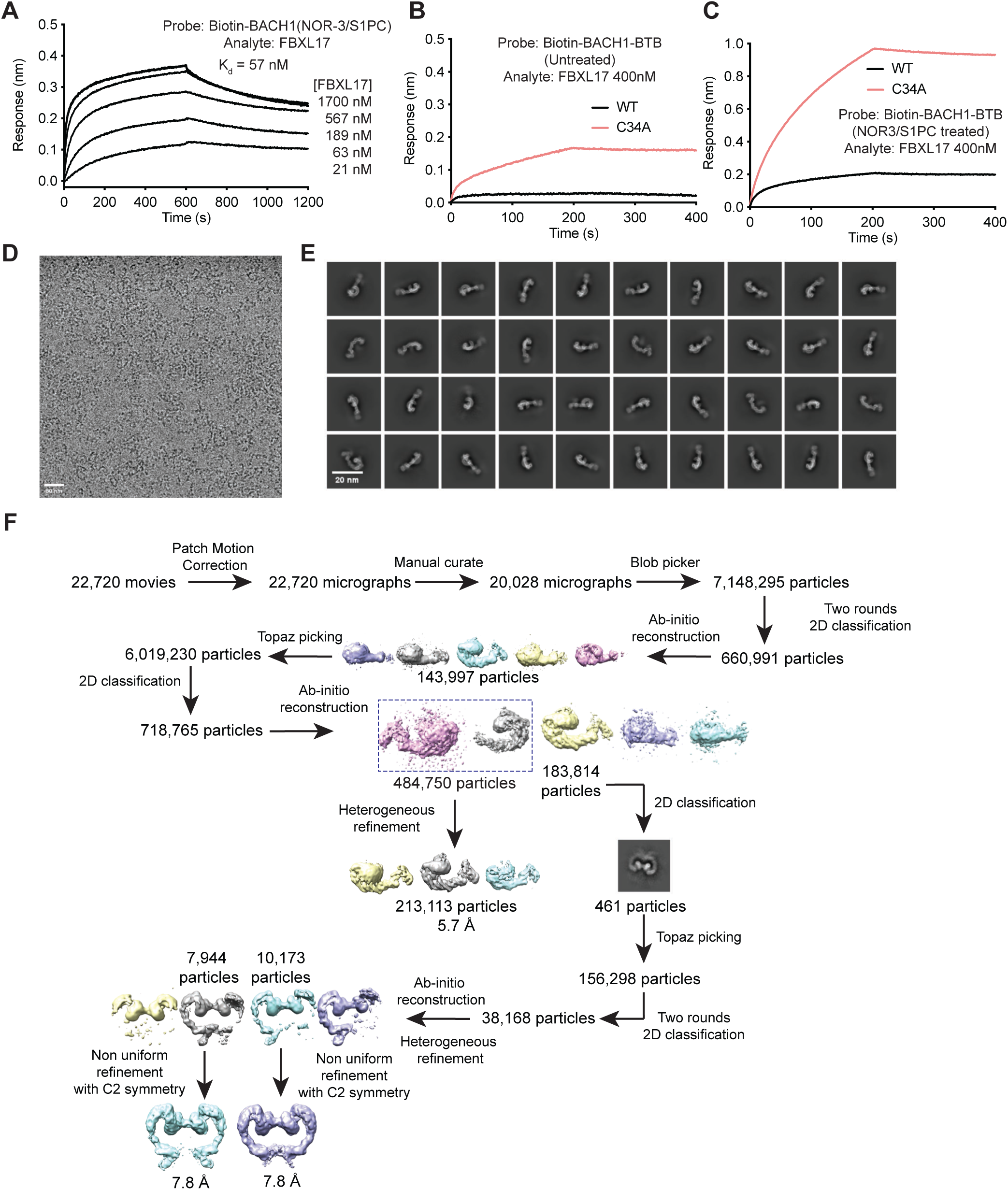
BLI analysis of FBXL17-BACH1-BTB interactions and the schematic workflow of single particle reconstruction of the SCF^FBXL17-BACH1^complex. A. BLI measurements of the binding between FBXL17 and NOR3-S1PC-treated BACH1-BTB with a 10-minutes association step. Kd, dissociation constant. **B. & C.** BLI measurements of the interaction between FBXL17 and BACH1-BTB (wild type and C34A mutant) untreated or treated with NOR3-S1PC. In the absence of compound treatment, C34A enhanced FBXL17 binding. This effect is exaggerated upon compound treatment. The C34 residue, therefore, is not required for S-nitrosylation. **D.** A representative cryo-EM micrograph for the sample containing a mixture of SCF^FBXL17^ and BACH1-BTB treated with NOR3-S1PC. **E.** Typical 2D averages. **F.** The flowchart of single particle analysis of the sample containing a mixture of SCF^FBXL17^ and BACH1-BTB treated with NOR3-S1PC.

**Figure S5.**
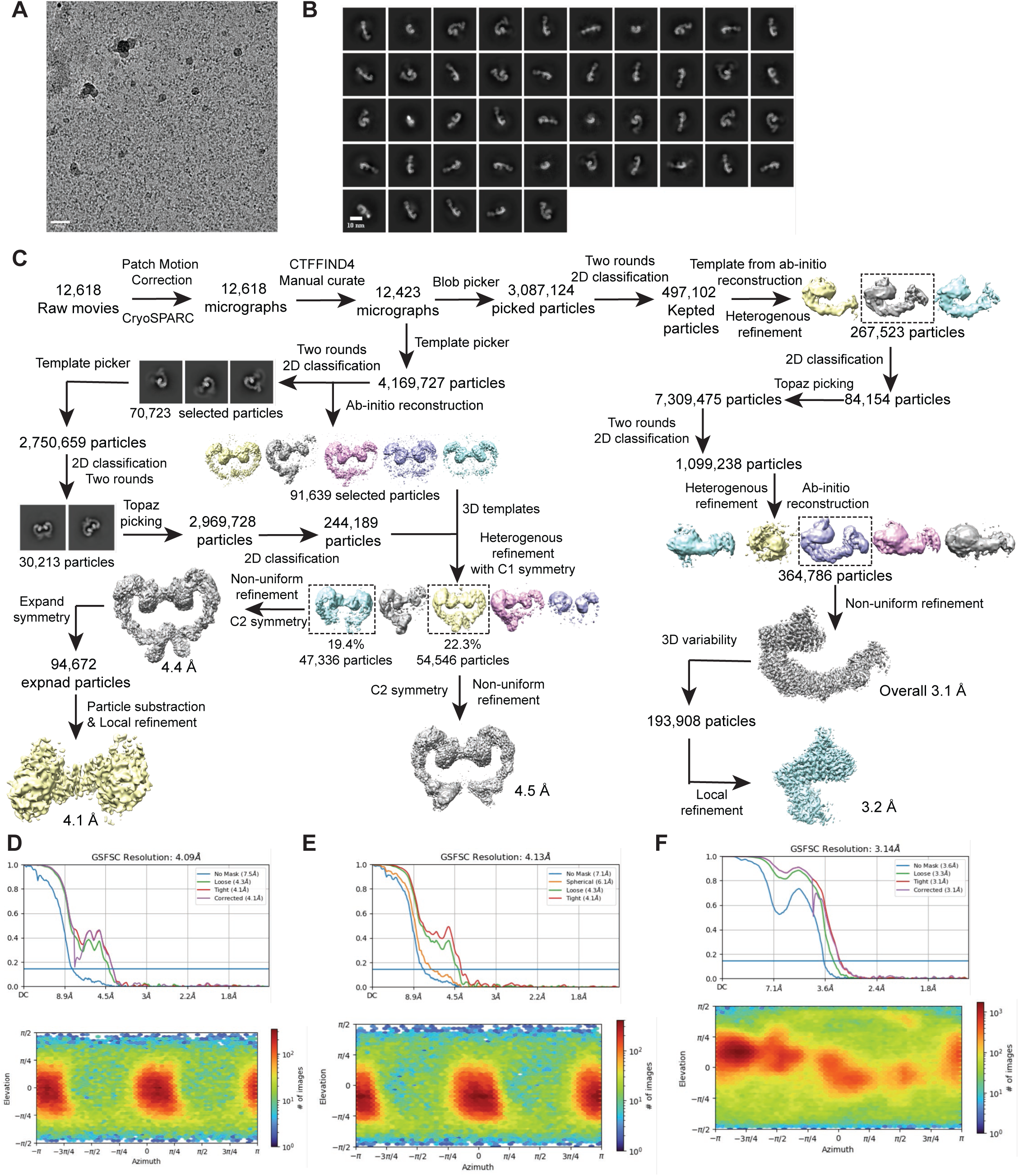
Cryo-EM single particle analysis workflow of wild type SCF^FBXL17-BACH1^. A. A representative cryo-EM micrograph. **B.** Typical 2D averages revealing the most populated monomeric complex. **C.** The flowchart of single particle analysis of the sample containing a mixture of SCF^FBXL17^ and BACH1-BTB. **D.** The particle angular distribution and FSC curves of dSCF^FBXL17-BACH1-I^ with local refinement. **E**. The particle angular distribution and FSC curves of dSCF^FBXL17-BACH1-II^. **F.** The angular distribution and FSC curves of the monomeric SCF^FBXL17-BACH1^ complex.

**Table S1.** Cryo-EM data collection, refinement, and validation statistics.

|  | **SCF<sup>FBXO22</sup>_**<br>**BACH1-BTB** | **SCF<sup>FBXL17</sup>_**<br>**BACH1-BTB**<br>**Monomer** | **SCF<sup>FBXL17</sup>_**<br>**BACH1-BTB**<br>**Dimer** |
| --- | --- | --- | --- |
| **Data collection and processing** |  |  |  |
| Magnification | 105,000 | 165,000 | 165,000 |
| Voltage (kV) | 300 | 300 | 200 |
| Electron exposure (e<sup>-</sup>/Å<sup>2</sup>) | 59 | 60 | 60 |
| Defocus range (μm) | -1.8~-3.5 | -0.8~-2.5 | -0.8~-2.5 |
| Pixel size (Å) | 0.84 | 0.743 | 0.743 |
| Symmetry imposed | C1 | C1 | C2 |
| Initial particle images (no.) | 2,888,035 | 3,087,124 | 2,969,728 |
| Final particle images (no.) | 413,523 | 364,786 | 71,155 |
| Map resolution (Å) | 3.9 | 3.1 | 4.1 |
| FSC threshold | 0.143 | 0.143 | 0.143 |
| Map resolution range (Å) | 3.9~6.5 | 3.1~6.5 | 4.1~7.0 |
| **Refinement** |  |  |  |
| Initial model | AlphaFold 2 | 6WCQ/1LDJ | SCF-FBXL17-BACH1-BTB |
| Map sharpening *B* factor (Å<sup>2</sup>) | -243.4 | -99.6 | -145.5 |
| Model composition |  |  |  |
| Non-hydrogen atoms | 9,822 | 8,798 | 23,028 |
| Protein residues | 1,232 | 1,101 | 2861 |
| Ligands | - | - | - |
| Zn<sup>2+</sup> | - | - | 6 |
| *B*-factors (Å<sup>2</sup>) |  |  |  |
| Protein | 85.24 | 28.12 | 17.28 |
| Ligand | - | - | 15.57 |
| R.m.s. deviations |  |  |  |
| Bond lengths (Å) | 0.005 | 0.004 | 0.0044 |
| Bond angles (°) | 1.078 | 0.809 | 0.823 |
| Validation |  |  |  |
| MolProbity score | 2.25 | 2.28 | 2.19 |
| Clashscore | 13.68 | 10.25 | 12.59 |
| Poor rotamers (%) | 0 | 0 | 0.1 |
| Ramachandran Plot |  |  |  |
| Favored (%) | 87.73 | 86.2 | 90.21 |
| Allowed (%) | 11.53 | 13.16 | 9.61 |
| Disallowed (%) | 0.74 | 0.64 | 0.18 |

## Data availability

The atomic model of FBXO22-BACH1-BTB from *Homo sapiens* and the cryo-EM maps were deposited to the PDB (https://www.rcsb.org) with accession number 8UA3 and to the EMDB (https://www.ebi.ac.uk/emdb) with the accession number EMD-42049. The coordinates of SCF^FBXO22^-BACH1-BTB and cryo-EM maps from the same dataset of FBXO22-BACH1-BTB were also deposited to the PDB (https://www.rcsb.org) with accession number 8UA6 and to the EMDB (https://www.ebi.ac.uk/emdb) with the accession number EMD-42051. The atomic coordinates and EM maps of wild-type BACH1-BTB bound FBXL17 were deposited to PDB with accession number 8UAH and to EMDB with accession number EMD-42064, and the atomic model and EM map of overall SCF^FBXL17^-BACH1-BTB were also deposit to PDB and EMDB with accession number 8UBT, EMD-42102 respectively. The atomic coordinates and EM maps of dimeric SCF^FBXL17^-BACH1-BTB and FBXL17LRR-BACH1-BTB were deposited to PDB and EMDB with accession numbers 8UBU and EMD-42105, 8UBV, and EMD-42106, respectively. The EM map of dimeric SCF^FBXL17^-BACH1-BTB open conformation was deposited to EMDB with an accession number EMD-42115.

